# Genetic and immune determinants of *E. coli* liver abscess formation

**DOI:** 10.1101/2023.06.11.543319

**Authors:** Karthik Hullahalli, Katherine G. Dailey, Yuko Hasegawa, Masataka Suzuki, Hailong Zhang, David W. Threadgill, Matthew K. Waldor

## Abstract

Systemic infections can yield distinct outcomes in different tissues. In mice, intravenous inoculation of *E*. *coli* leads to bacterial replication within liver abscesses while other organs such as the spleen largely clear the pathogen. Abscesses are macroscopic necrotic regions that comprise the vast majority of the bacterial burden in the animal, yet little is known about the processes underlying their formation. Here, we characterize *E. coli* liver abscesses and identify host determinants of abscess susceptibility. Spatial transcriptomics revealed that liver abscesses are associated with heterogenous immune cell clusters comprised of macrophages, neutrophils, dendritic cells, innate lymphoid cells, and T-cells that surround necrotic regions of the liver. Susceptibility to liver abscesses is heightened in the C57BL/6 lineage, particularly in C57BL/6N females. Backcross analyses demonstrated that abscess susceptibility is a polygenic trait inherited in a sex-dependent manner without direct linkage to sex chromosomes. As early as one day post infection, the magnitude of *E. coli* replication in the liver distinguishes abscess-susceptible and abscess-resistant strains of mice, suggesting that the immune pathways that regulate abscess formation are induced within hours. We characterized the early hepatic response with single-cell RNA sequencing and found that mice with reduced activation of early inflammatory responses, such as those lacking the LPS receptor TLR4, are resistant to abscess formation. Experiments with barcoded *E. coli* revealed that TLR4 mediates a tradeoff between abscess formation and bacterial clearance. Together, our findings define hallmarks of *E. coli* liver abscess formation and suggest that hyperactivation of the hepatic innate immune response drives liver abscess susceptibility.

**Importance:** Animal models of disseminating bacterial infections are critical for developing therapeutic interventions. Following systemic dissemination in mice, *E. coli* undergo dramatic replication within abscesses in the liver but not in other organs. Although liver abscesses are the largest reservoir of bacteria within the animal, the processes that lead to abscess formation are not known. Here, we characterize *E. coli* liver abscess formation and identify several determinants of abscess susceptibility, including sex, mouse genotype, and innate immune factors. By combining spatial and single-cell transcriptomics with genetic and phenotypic analyses, we delineate critical host pathways that underlie abscess formation. Our findings define several avenues for future studies to unravel how abscess susceptibility determinants interact to modulate clearance of systemic infections and govern tissue-specific bacterial replication.

## Introduction

Bloodstream infections are a leading cause of human mortality (1). Although bacteria routinely breach epithelial barriers and enter systemic circulation, most of these events do not cause disease, in large part because innate immune cells within the liver, spleen, and other organs sequester and kill circulating bacteria (2). However, many microorganisms encode factors that facilitate evasion of or resistance to these host defenses. Gram-negative species pose an especially challenging threat to the healthcare system due to the continued emergence of antimicrobial resistance (3).

The Gram-negative bacterium *Escherichia coli* is among the leading causes of human bloodstream infections (4). Due to the systemic nature of these infections, resolution of infection requires most organs in the host to mount an immune response, and these responses vary across tissues. Consequently, systemic infections caused by Extraintestinal Pathogenic *E. coli* (ExPEC) can manifest a wide range of tissue-specific clinical syndromes in which bacterial factors, such as pili and siderophores, enable the pathogen to counteract host defenses and survive and replicate (5–7).

Ultimately, the interplay between these pathogen factors and host defenses leads to tissue-specific pathology (8). Deciphering why some tissues are permissive to pathogen growth while others are restrictive is critical for understanding the mechanistic underpinnings of infection outcomes across host tissues. Existing models of systemic ExPEC infection yield either rapid sepsis and death within hours (9, 10) or clearance of bacteria from the animal (11, 12). An animal model that lies in between these two extremes, where bacteria replicate and survive within the host for extended time periods, would deepen our understanding of pathogen and host factors that influence the outcome of extraintestinal *E. coli* infections.

We previously observed that mice inoculated intravenously with ExPEC developed visibly apparent abscesses specifically in the liver (15). By using a library of bacteria that possessed ∼1000 unique DNA barcodes at a neutral locus and the STAMPR computational pipeline (13), we found that abscesses coincide with the expansion of ∼10 clones that replicate to ∼10^7^ colony forming units (CFU) (14). Although abscesses in the liver represent the predominant site of *E. coli* replication in the animal, the mechanisms that underlie *E. coli* abscess formation are unknown. In general, animal models of Gram-negative liver infections have received little attention. Since *E. coli* is a leading cause of human liver abscesses and abscesses are often fatal if left untreated (15), a tractable animal model is valuable for expanding understanding of tissue specific immune responses and for the development of therapeutic interventions.

In this study, we characterize the cellular composition, genetics, kinetics, and immunology of *E. coli*-induced liver abscesses in mice. Liver abscesses are dependent on mouse genotype and are inherited in a sex-dependent manner without direct linkage to sex chromosomes. Although abscesses require several days to fully develop, the pathways that confer susceptibility to abscess formation are engaged within hours, the timescale in which massive numbers of innate immune cells are recruited to the liver and proinflammatory cytokines are induced. Mice that are resistant to abscess formation are defective for both Gr1+ inflammatory cell recruitment and proinflammatory cytokine production in the hours following inoculation. These defects are phenocopied in mice lacking the LPS receptor TLR4, which are comparably resistant to abscess formation. However, in the absence of TLR4, fewer *E. coli* are eliminated by host restriction processes, suggesting that TLR4 governs a tradeoff between pathogen clearance and replication. We propose that *E. coli* liver abscesses result when tissue damage from inflammation provides a niche for pathogen replication. Taken together, our findings reveal important characteristics of a mouse model for *E. coli*-induced liver abscesses and establish a tractable platform to investigate tissue-specific innate immunity.

## Results

### Phenotypic characterization of *E. coli-*induced liver abscesses

Female C57BL/6J (B6J) mice were inoculated intravenously (IV) with 5×10^6^ CFU of a barcoded *E. coli* library (CHS7-STAMP, derived from extraintestinal pathogenic strain CFT073 (16)) and liver bacterial burden was enumerated at 5 days post inoculation (dpi). Consistent with our previous study (14), we found that approximately half of the mice developed visible liver abscesses (approximately 0.5 - 2 mm^2^) with correspondingly very high bacterial burdens (Figure 1); animals with abscesses had 100,000 times greater CFU than those that did not. Occasionally, animals with very low CFU had very small white lesions. In this study, abscesses are defined as the co-occurrence of visible white lesions and a hepatic CFU burden of at least 10^4^. Hematoxylin and Eosin (H&E) staining revealed that the abscess core primarily consists of necrotic hepatocytes surrounded by mixed inflammatory cells resembling macrophages and neutrophils (Figure 1, Figure S1A). Despite the apparent tissue damage (Figure 1, Figure S1A), serum levels of alanine aminotransferase (ALT), which is released from damaged hepatocytes (17), were similar in animals that possessed or lacked abscesses (Figure S1B).

**Figure 1.**
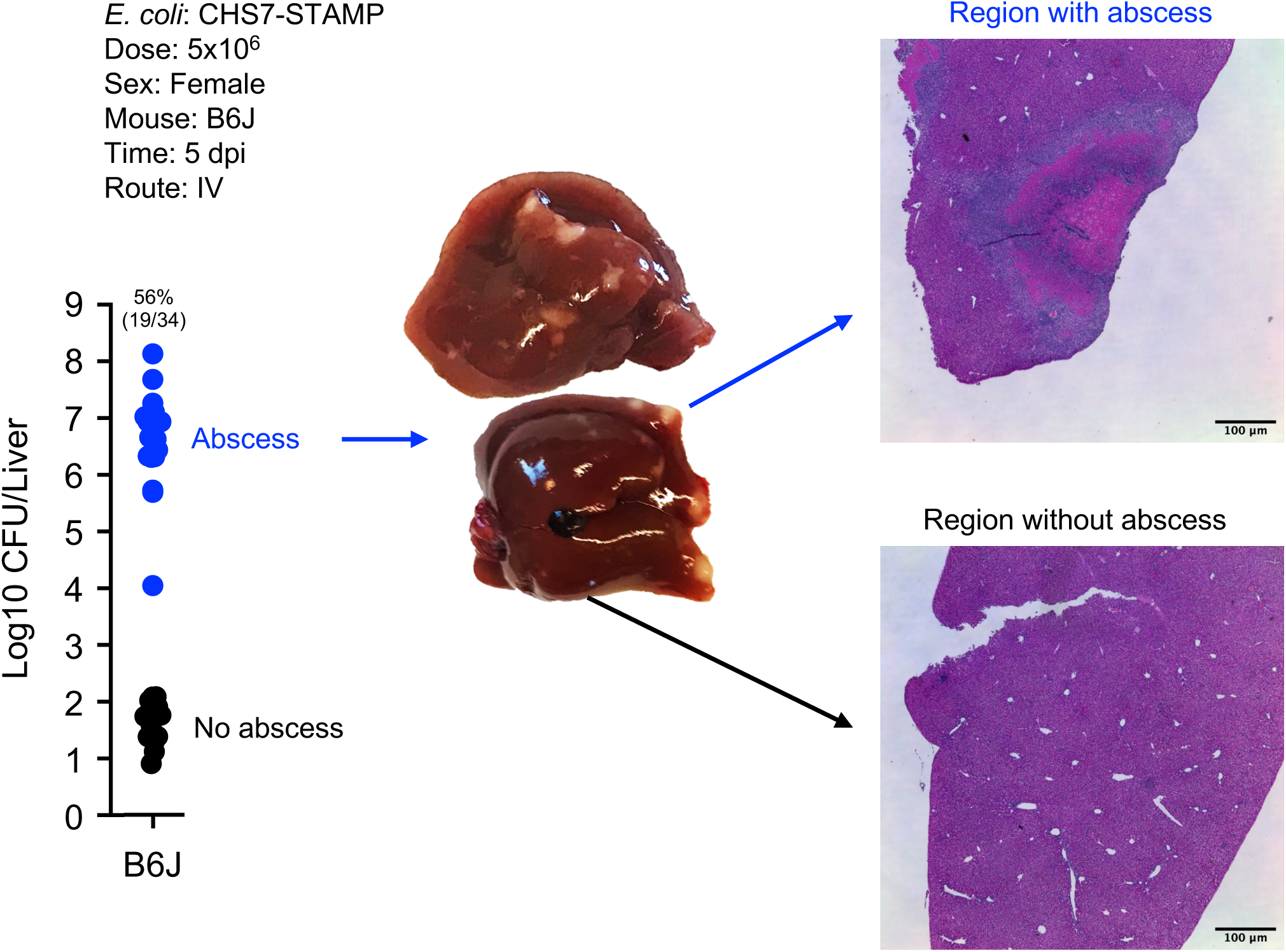
Intravenous inoculation of *E. coli* induces liver abscesses in mice. *E. coli* hepatic CFU burden in B6J female mice 5 dpi. Approximately half of animals form abscesses (blue), which are associated with marked bacterial replication; livers from animals that do not develop abscesses contain relatively few *E. coli* (black). Images of livers containing abscesses are shown as well as H&E staining. Additional images are shown in Figure S1.

Unlike B6J females, BALB/cJ, CBA/J, and C3H/HeJ females were entirely resistant to abscess formation (Figure 2A). Surprisingly, female C57BL/6NJ (B6N) mice developed abscesses at even higher frequencies than B6J females. Increased abscess frequency in B6N relative to B6J was also observed at a 10-fold lower inoculum size, where B6J mice do not develop abscesses (Figure 2B). B6N and B6J diverged from the ancestral C57BL/6 strain in 1951 following their transfer to the National Institutes of Health (B6N) from Jackson Labs (B6J) and differ by ∼10,000 SNPs as of 2013 (18). These data suggest that the allele(s) conferring abscess susceptibility is present in the C57BL/6 lineage, and one or more mutations have occurred within this lineage that further distinguish B6N and B6J.

**Figure 2.**
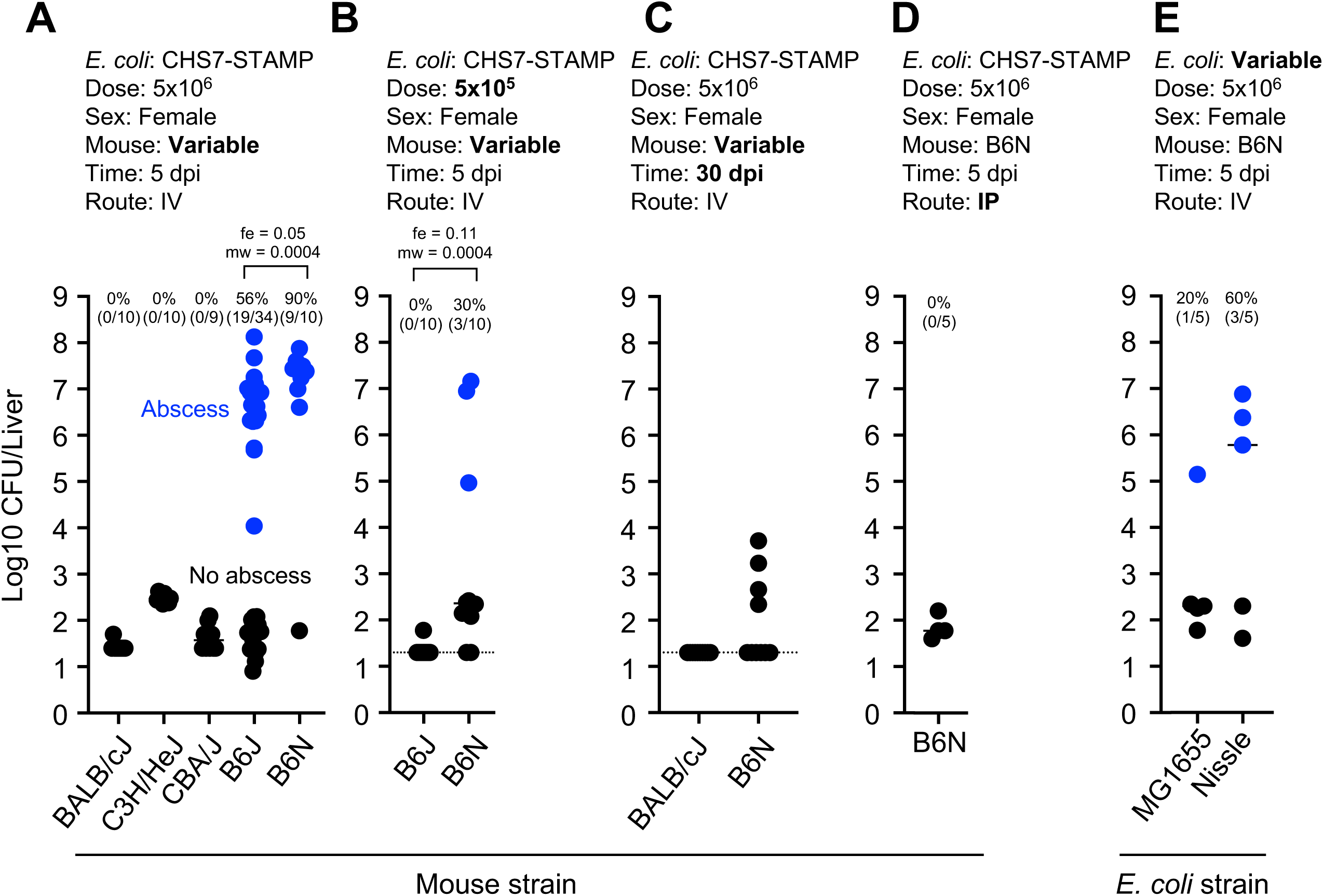
Susceptibility to *E. coli* liver abscess varies in different inbred mouse strains. Blue points represent animals that developed abscesses. Experimental parameters are included above each plot, and bolded text highlights key variable parameters. Abscess frequencies and exact numbers of animals are shown above each group. P values are derived from one tailed Mann Whitney U tests (mw) and Fisher Exact Tests (fe). A) Abscesses are specific to the C57Bl/6 lineage. These data were used to define >10,000 CFU and visible abscess formation as criteria for defining abscesses. B) Differences in infection outcome between B6J and B6N females were also apparent at a lower dose. C) Abscesses are cleared by 30 dpi. D) Abscesses do not form in B6N females following IP injection. E) Commensal *E. coli* MG1655 and Nissle can also stimulate abscess formation. Dotted lines in B and C represent limits of detection.

Abscess susceptibility did not correlate with other clinical outcomes. B6N (susceptible) females lost weight at 1 dpi but then their weight remained stable at 5 dpi, when abscesses have fully formed (Figure S1C). B6N females also survived up to 30 dpi, at which point abscesses were no longer present and there were <10^4^ CFU in livers (Figure 2C). Therefore, *E. coli* liver abscesses do not cause mortality and are eventually cleared in mice. No *E. coli* CFU were detectable in female BALB/cJ (resistant) at 30 dpi. Abscesses did not occur in B6N females when the route of inoculation was changed from IV to intraperitoneal injection, suggesting that immediate pathogen capture by the liver may be important for abscess formation (Figure 2D). Abscesses also formed in B6N females following IV inoculation of nonpathogenic *E. coli* strain Nissle (19), and to a lesser extent with laboratory strain MG1655, suggesting that abscess induction is not a unique property of the bacterial strain used in this study (Figure 2E).

To gain further insight into the identity and function of the immune cells that surround the necrotic zone within abscesses, we performed spatial transcriptomics with MERFISH (Multiplexed Error Robust Fluorescence In Situ Hybridization) a probe-based hybridization technique (20), using the MERSCOPE instrument (Figure 3, Figure S2-S5). MERFISH enables simultaneous identification of hundreds of user-specified RNA molecules *in situ.* We selected transcripts corresponding to specific cell types identified in the Liver Cell Atlas (21). In uninfected B6J animals, hepatocyte zonation markers (*Cyp2e1* and *Cyp2f2*) clearly demarcated differential expression across hepatocytes, indicating that this approach is useful for analysis of liver tissue. By 3 dpi, substantially reduced RNA signal was observed within abscess cores, which lacked hepatocyte zonation markers presumably due to local necrosis and RNA degradation (Figure 3, Figure S2-S5). On the border of and within 5 dpi abscesses, enrichment of transcripts corresponding to migratory dendritic cells (*Cacnb3*), neutrophils (*S100a8/9*) and macrophages (*Adgre1*) were detected. Transcripts corresponding to T cells (*Cd4, Cd3g, Cd3e, Cd3d)* and NK cells/ILC1s (*Klrb1b)* cells were also detected in these locations but at lower abundances. The assembly of similar, smaller immune cell clusters was also seen associated with smaller zones of RNA degradation at 3 dpi. Some immune cell clusters lacked RNA degradation altogether and may represent resolved or early-stage abscesses (Figure S3). Importantly, these cell clusters were absent in uninfected mice (Figure 3, Figure S2) and were enriched for lysozyme (*Lyz2*), nitric oxide synthase (*Nos2*), and cytochrome b oxidase (*Cybb*), markers associated with inflammatory responses (Figure 3, Figure S2-S5). These results indicate that immune cell clusters associated with liver abscesses are heterogeneous and express many known inflammatory markers.

**Figure 3.**
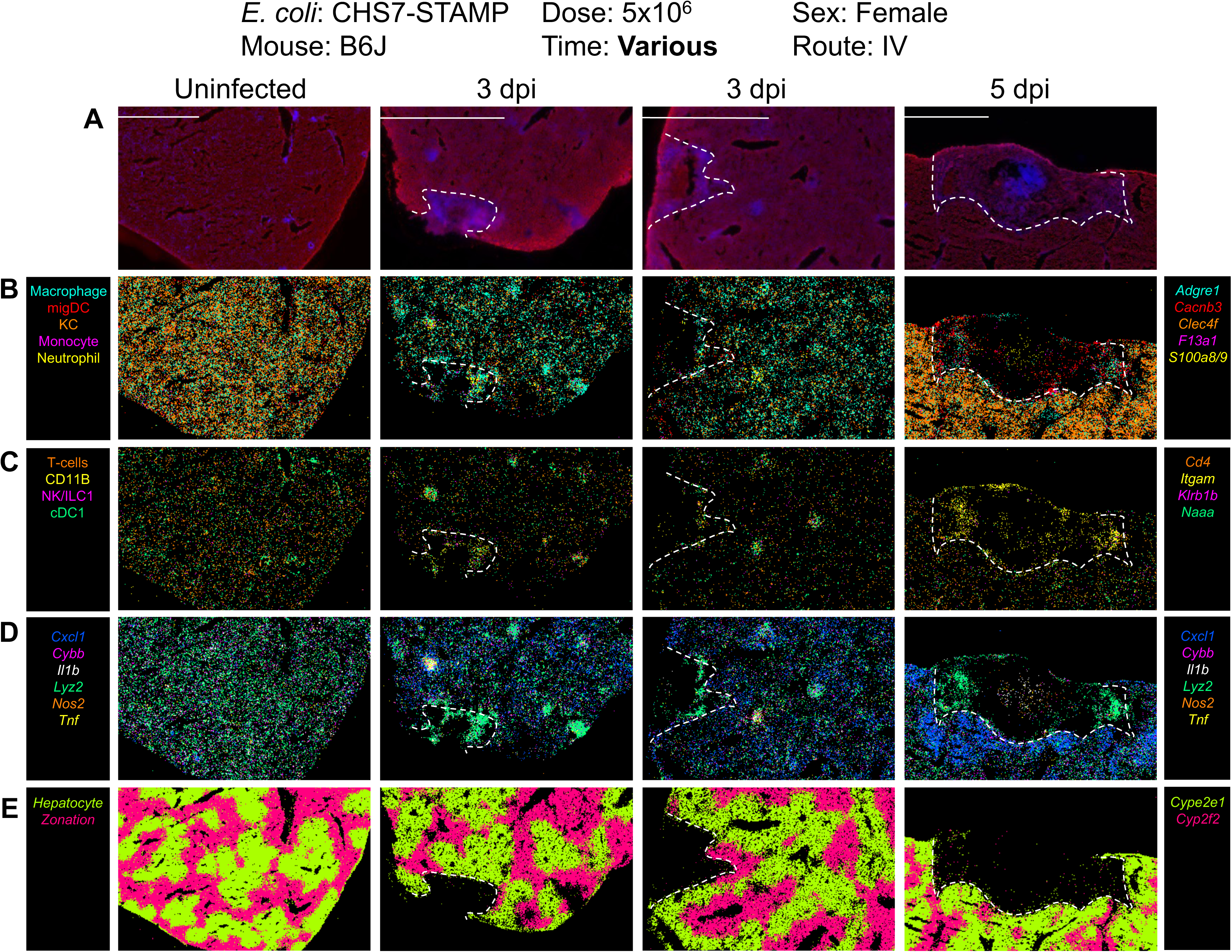
Spatial transcriptomic profiles of liver abscesses. MERSCOPE images of liver samples from uninfected, 3 dpi, and 5 dpi mice. Unmerged images are shown in Figure S2-S4 and quantification is shown in Figure S5. Dotted lines denote abscesses, which coincide with RNA degradation. Transcripts were selected from the liver cell atlas (21), which defines specific cell types associated with each transcript. A) Abscess boundary and DAPI staining. B) Macrophages (*Adgre1),* migratory dendritic cells (*Cacnb3*), Kupffer cells (*Clec4f*), monocytes (*F13a1*), and neutrophils (*S100a8/9*). C) T cells (*Cd4*, also *Cd3g*, *Cd3e*, and *Cd3b*, which were not marked in figure for clarity), NK/ILC1s (*Klrb1b*), and cDC1 (*Naaa*). *Itgam* (CD11b) is a marker for multiple leukocyte subsets, including macrophages, dendritic cells, granulocytes, and NK cells. D) Markers of inflammation. E) Markers of liver zonation.

### Abscess susceptibility is a polygenic trait with sex-influenced inheritance

To begin to identify host factors that regulate liver abscess formation, we first determined the inheritance pattern of the abscess susceptibility trait. Importantly, both sexes of BALB/cJ mice were resistant to abscesses, while both sexes of B6J mice were sensitive, with slightly heightened sensitivity in B6J males (Figure 4A). The F1 offspring of female BALB/cJ (resistant) and male B6J mice (sensitive), known as CB6F1/J, were challenged intravenously with *E. coli* and CFU burden and the frequency of liver abscesses were assessed 5 dpi. Surprisingly, only F1 female mice inherited the abscess susceptibility trait, while F1 males were resistant (Figure 4B). Since male CB6F1/J mice lack a B6J X-chromosome, and female CB6F1/J mice have 1 copy each of the BALB/cJ and B6J X-chromosome, these data initially raised the possibility that abscess susceptibility is X-linked in B6J. In this scenario, the B6J X-chromosome possesses an abscess susceptibility allele, and CB6F1/J males lack this allele. To experimentally test if an abscess susceptibility allele is X-linked, we crossed female B6J mice with male BALB/cJ mice (reverse sexes from previous cross). In the F1 offspring (known as B6CF1), males possess a B6J X-chromosome, whereas females possess both BALB/cJ and B6J alleles. Surprisingly, B6CF1 males and females phenocopied CB6F1/J mice; males were resistant, and females were susceptible (Figure 4B). These data reveal that abscess susceptibility is not sex-chromosome linked in BALB/cJ x B6J F1 animals but is influenced by sex. However, the influence of sex on susceptibility seems to be modified by genetic background. We found that inbred B6N males were more resistant to abscess formation than B6N females (Figure 4A). Given the differences in abscess frequency between B6J and B6N mice, we assessed whether the inheritance pattern in BALB/cJ X B6N F1 animals was distinct from that observed in BALB/cJ X B6J F1 mice. However, we again found that F1 females were partially sensitive, and males were resistant regardless of the sex of the parents, similar to the F1 offspring of B6J and BALB/cJ (Figure 4C).

**Figure 4.**
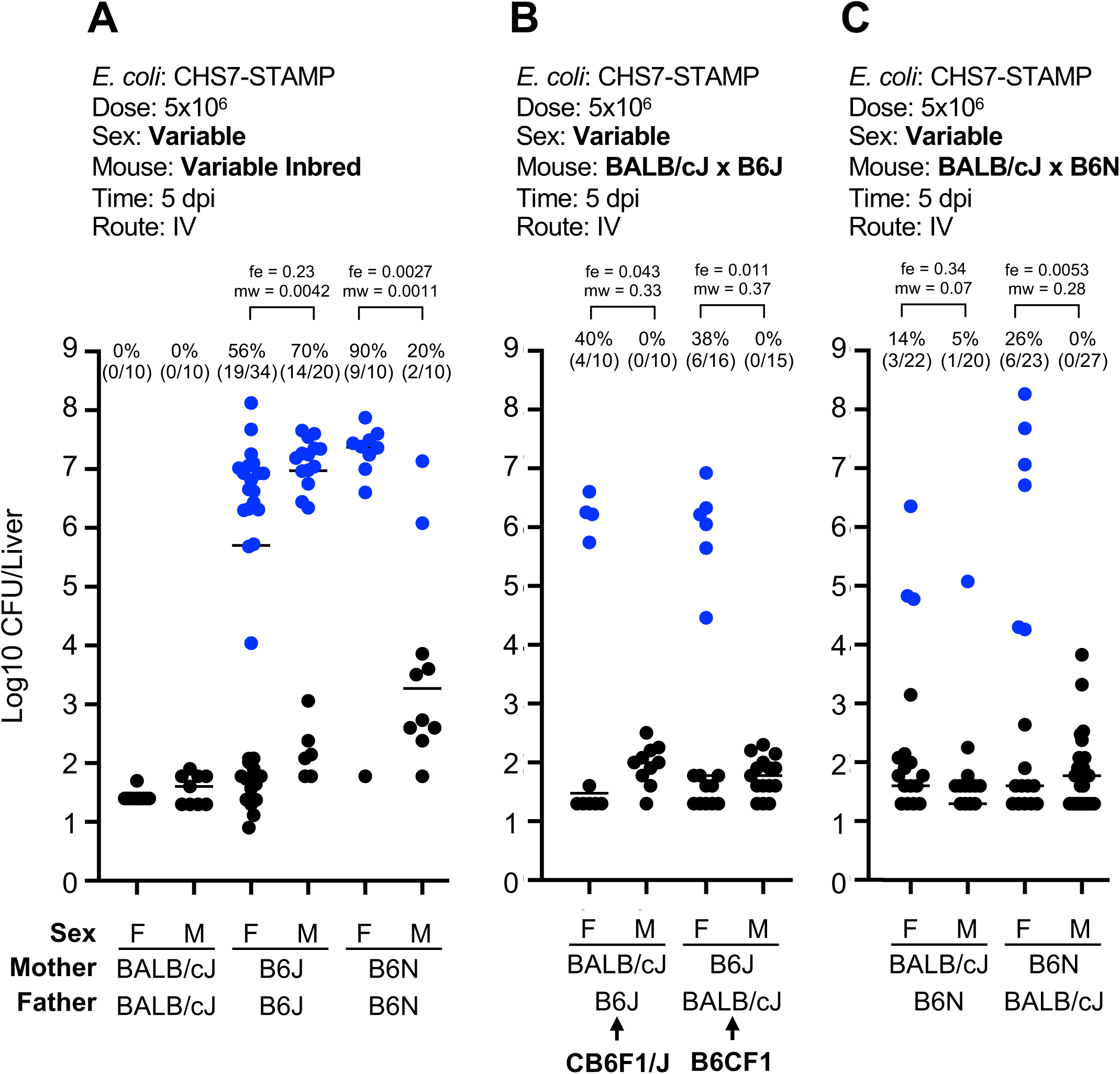
Abscesses are sex-linked in B6N and F1 heterozygous mice. Blue points represent animals that developed abscesses. Experimental parameters are included above each plot and bolded text highlights key variable parameters. Abscess frequencies and exact numbers of animals are included above each group. P values are derived from one tailed Mann Whitney U tests (mw) and Fisher Exact tests (fe). A) Female B6N mice are more susceptible to abscess formation than male B6N. In contrast, male B6J trend towards slightly increased abscess formation than females. Both BALB/cJ males and females are resistant to abscess formation. B) Abscess susceptibility is inherited only by females from crosses between B6J and BALB/cJ regardless of the sex of the parents. C) Same as B) but using B6N mice instead of B6J

These results cannot be explained by maternal inheritance of mitochondria; CB6F1/J males and B6CF1 males both differ in mitochondrial alleles but possess identical phenotypes. Furthermore, abscess susceptibility is not directly conferred by the Y-chromosome, since females are generally more sensitive, except in inbred B6J. The BALB/cJ Y-chromosome is also unlikely to possess a unique, abscess-inhibitory factor, since male F1 offspring are all resistant to abscess formation regardless of Y-chromosome alleles. Collectively, these results support a model where abscess susceptibility is inherited in a recessive manner in males and in an incomplete dominant manner in females, but is not directly linked to the X, Y, or mitochondrial chromosomes.

With the expectation that the abscess susceptibility trait is autosomal, we carried out backcrosses between B6J males and CB6F1/J females, generating N1 backcross offspring (Figure S6A). All N1s are identical for mitochondrial (all BALB/cJ from maternal grandmother) and Y chromosomes (all B6J from father). Among the autosomes, these N1s are ∼50% heterozygous and ∼50% homozygous for B6J alleles at random loci. Since abscesses are incomplete dominant in females, all N1 females should have a ∼50% likelihood of developing abscesses. However, since abscesses are inherited recessively in males, only male mice that are homozygous for the causal B6J allele should be susceptible (∼70% likelihood) to abscesses. Thus, we proceeded only with male N1 mice. 153 male N1 mice were analyzed with a genotyping array consisting of ∼3000 SNPs that distinguish BALB/cJ and B6J alleles to map the heterozygosity, homozygosity (B6J), or hemizygosity (for X-chromosome) of the N1 genomes. At 8-10 weeks of age, male N1s were infected and abscess frequency and CFU burden were scored at 5 dpi. Importantly, 44% of mice developed abscesses, confirming that introducing B6J alleles into an otherwise resistant background (heterozygous CB6F1/J males) also reintroduces abscess susceptibility (Figure S6B).

At every SNP, we calculated abscess frequencies of homozygous (for B6J) mice relative to abscess frequencies of heterozygous mice, expecting that abscesses would be more likely to form in mice homozygous at the causal allele relative to mice that are heterozygous for the causal allele. Given that N1s consist of mice with both brown and black coats, we verified that this strategy is effective by identifying the agouti locus (which governs coat color). We observed a clear signal for homozygosity in Chr. 2 at the location of the agouti locus in mice with black coats (Figure S6C). However, we observed no such signal when identifying homozygous loci associated with abscess formation (Figure S6D). Since this approach can identify a monogenic trait, we conclude that the abscess susceptibility trait is polygenic. Specifically, B6J males contain at least two loci that are, when homozygous, independently sufficient to confer abscess susceptibility. Abscess susceptibility in heterozygous F1 females suggests that these loci only require one copy to sensitize females to abscess formation.

### Bacterial replication and early hepatic responses following infection

The backcross experiments did not lead to the identification of a single genetic locus that distinguishes abscess-susceptible versus abscess-resistant mice. Therefore, we set out to identify phenotypes associated with abscess formation that may distinguish susceptible and resistant mouse strains. Identifying these phenotypes required further knowledge of the kinetics of abscess formation to facilitate distinguishing between pathways that cause abscesses and those that simply respond to the increased bacterial burden associated with abscess formation.

We examined whether female BALB/cJ (resistant), B6J (intermediate-susceptible), and B6N (hyper-susceptible) mice had phenotypically diverged as early as 1 dpi. Indeed, by 1 dpi, total burden was low in BALB/cJ, intermediate in B6J, and high in B6N (Figure 5A). Furthermore, bacterial burden correlated with gross appearance of the liver; even at 1dpi there were prominent white lesions in the livers of B6N female mice that were less abundant in B6J and absent in BALB/cJ animals (Figure 5B). Sequencing the barcode loci to measure the abundance of individual clones confirmed that these differences in CFU correlated with increased replication of a small number of clones, the bacterial hallmark of abscess formation. At 1 dpi BALB/cJ livers lacked replicating clones, while B6J had 1-2 replicating clones, and B6N had ∼40 replicating clones (Figure 5C, Figure S7). Bacteria within abscess-susceptible mice therefore have a higher likelihood of undergoing replication early after inoculation, which presumably drives abscess formation. Collectively, these data suggest that the innate immune pathways that underlie abscess-sensitive and resistant phenotypes likely diverge within the first day, when early signs of abscess formation are already apparent, both as visible lesions in the liver and as replication of *E. coli* clones.

**Figure 5.**
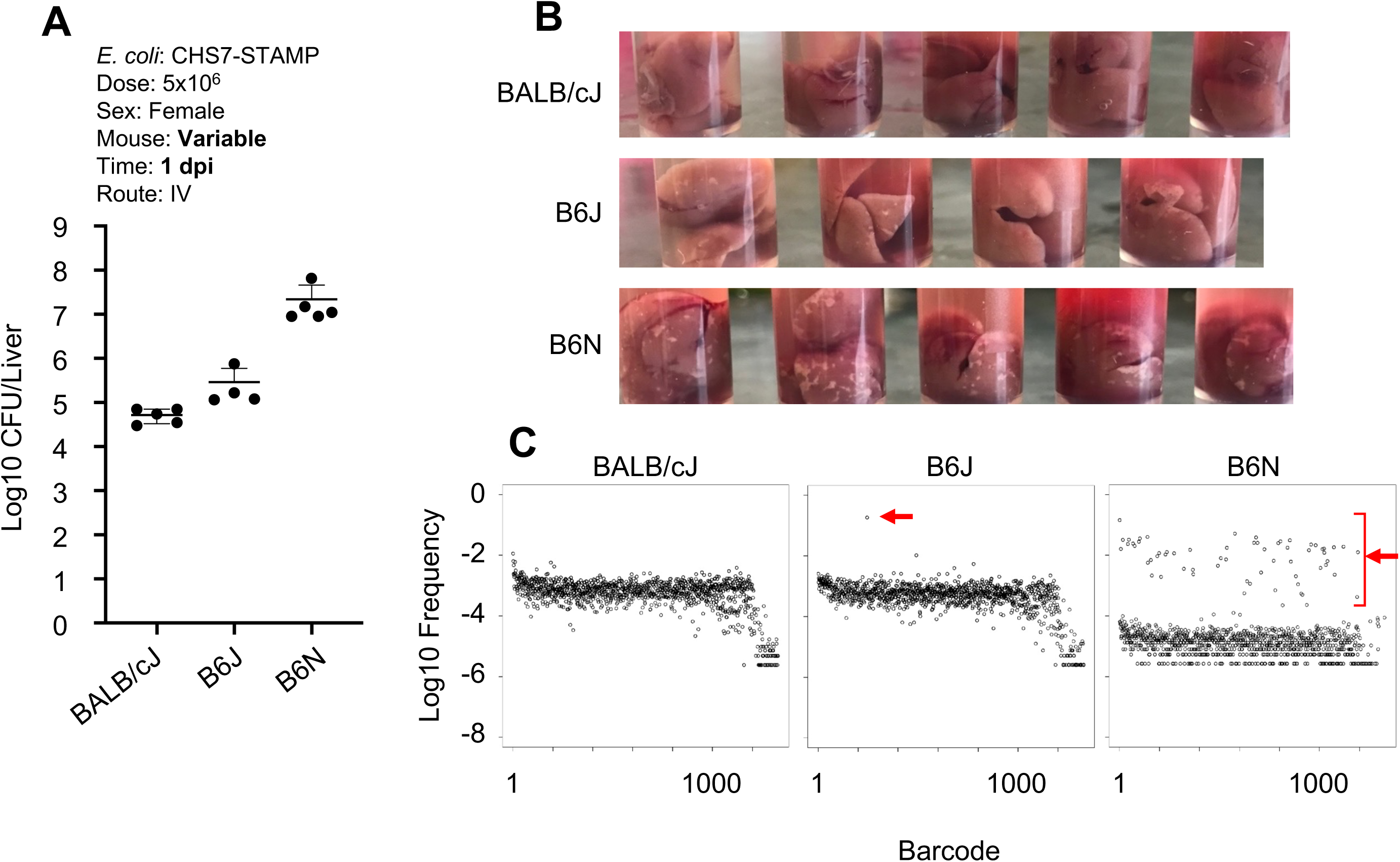
Abscess susceptibility correlates with phenotypes at 1 dpi. A) Total liver *E. coli* CFU at 1 dpi in BALB/cJ, B6J, and B6N strains. B) White lesions in the livers of BALB/cJ, B6J, and B6N mice at 1 dpi C) Barcode distributions from 1 dpi mice. The X-axis is an arbitrary designation for barcode identity, and the Y-axis represents the relative frequency of each barcode. Red arrows denote clones that replicated. Additional replicate mice are shown in Figure S7.

We performed single cell RNA sequencing of liver CD45+ immune cells 4 hours post infection (hpi) to characterize the hepatic immune pathways activated early following infection, but prior to gross liver damage and *E. coli* replication (14). Because abscesses are dose-dependent in B6J mice (22), we inoculated mice at a range of inoculum sizes, from 0 CFU to 1×10^7^ CFU. UMAP clustering revealed a marked dose-dependent expansion of clusters (1, 3, 2, and 11) that expressed markers corresponding to primarily macrophages and neutrophils (Figure 6AB, Figure S8). The large magnitude of the increase in the abundance of these cell types at this early time strongly suggests that these cells infiltrated into the liver from the blood rather than expanded in situ. These infiltrating innate immune cells share similar gene expression patterns, including the expression of *S100a8/9*, *Lcn2*, and *Il1b,* and cluster together in UMAP space (Figure S8). Other cell types, including B cells, T cells, NK cells, and dendritic cells, did not change in relative abundance (Figure 6CD) but were also responsive to infection. These included the dose-dependent production of interferon gamma (*Ifng*) by NK/T cells, cytokine and chemokine production from T and B cells (*Cxcl1*, *Cxcl10*), and downregulation of growth factor signaling in endothelial cells (*Kdr*) (Figure S8). Taken together, these findings reveal that innate immune responses in the liver, which include macrophage and neutrophil infiltration and proinflammatory cytokine production, are induced prior to the replication of clones and visible liver damage that distinguish abscess-susceptible and resistant mouse strains. These inflammatory responses are correspondingly diminished at lower inoculum sizes where abscesses are less likely to develop.

**Figure 6.**
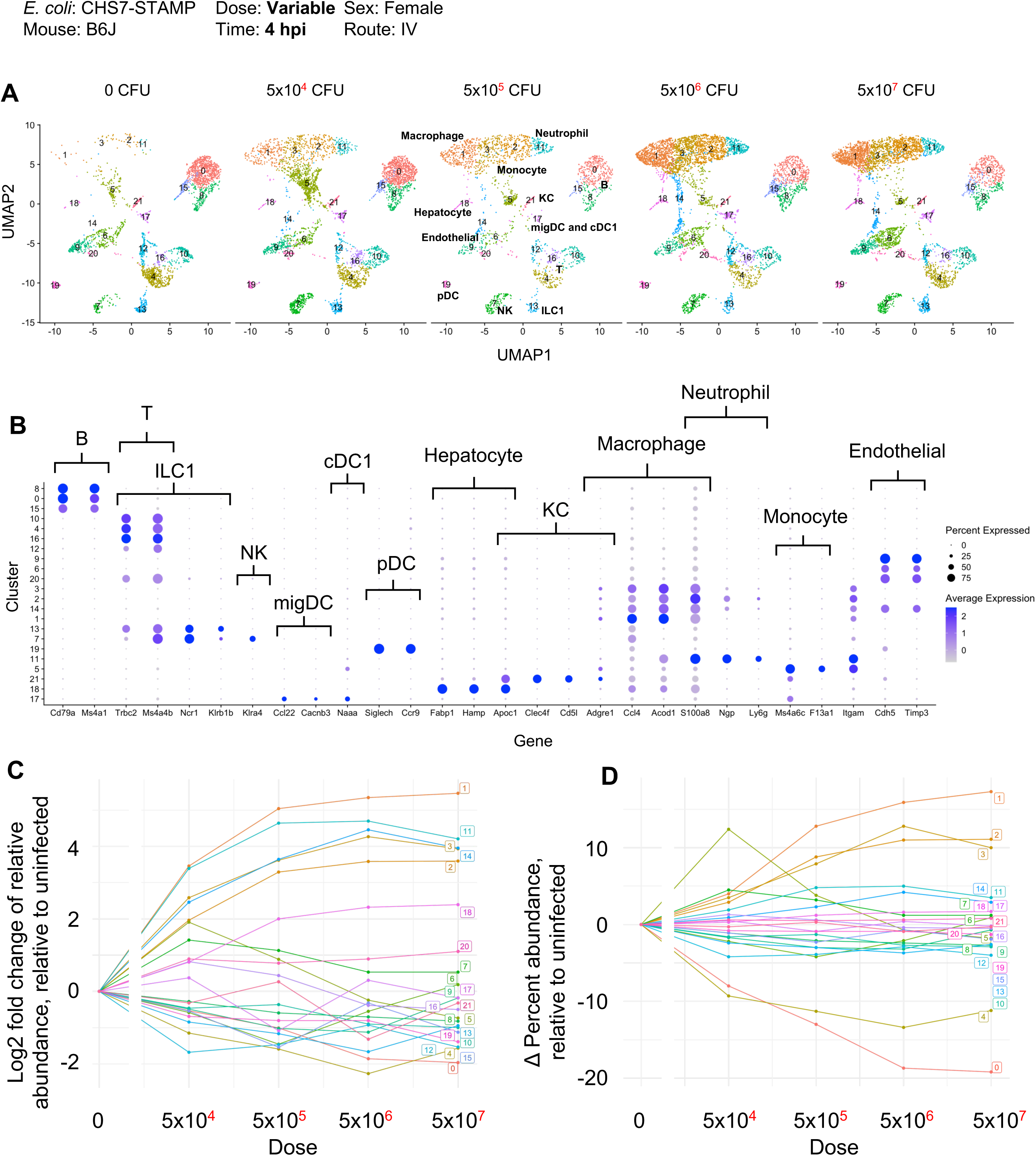
Single cell RNA sequencing of liver immune cells at 4hpi. A) UMAP plots from various inoculum sizes from CD45-sorted cells are shown. Infection results in a dose-dependent increase in the infiltration of clusters 1, 3, 2 and 11, corresponding to macrophages and neutrophils. B) Dot plots from normalized expression values (sctransform) at all doses of select genes used to classify clusters by cell type. C) Quantification of the fold change of relative abundance of each cluster, relative to uninfected. D) Same as C but displaying the absolute change in percentage abundance for each cluster, relative to uninfected.

### Abscess formation requires TLR4

To assess whether infiltrating immune cells directly contribute to abscess formation, we treated mice with an anti-Gr1 antibody, which depletes neutrophils and Ly6C^+^ monocytes and macrophages (23). However, Anti-Gr1 treatment resulted in 100% mortality by 2 dpi (Figure 7A) suggesting that at least some Gr1^+^ cell infiltration is required to control infection. We reasoned that a more subtle perturbation was necessary to elucidate the roles of early immune responses in the liver. Since many of the phenotypes observed at 4 hpi are likely induced by LPS stimulation via the LPS receptor Toll-like receptor 4 (TLR4) (24), we examined whether mice lacking TLR4 (in a B6J background) were resistant to abscess formation. Similar to McDonald et al, who found that TLR4^KO^ mice had reduced Gr1^+^ cell infiltration in the liver following IV LPS administration (25), Gr1^+^ cells were reduced in the livers of TLR4^KO^ mice after IV *E. coli* inoculation (Figure 7B). Further, serum levels of Cxcl1, Cxcl10, Il1β, and Tnfα, chemokines and cytokines that are downstream of TLR4 signaling, were reduced in TLR4^KO^ mice at 4 hpi. (Figure 7C). TLR4^KO^ females that were acquired from Jackson laboratories or bred in house failed to form abscesses, while TLR4^Het^ littermate controls were susceptible to abscess formation. However, CFU burden in TLR4^KO^ animals was higher compared to control animals that lacked abscesses (Figure 7D). These results indicate that abscess formation requires TLR4, consistent with the hypothesis that liver abscess susceptibility is driven by overactivation of the immune response.

**Figure 7.**
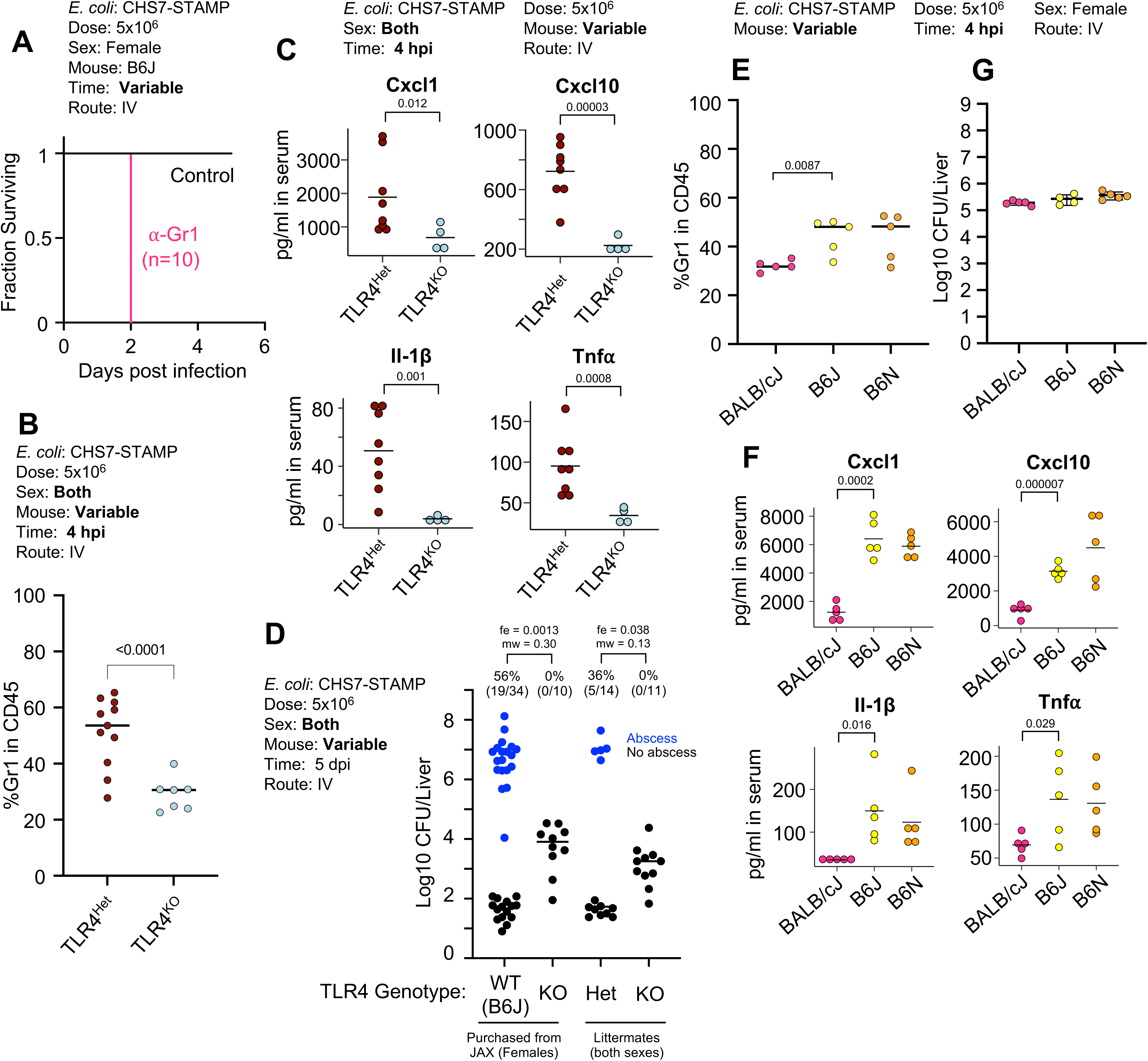
Abscess formation and immune responses to *E. coli* in TLR4^KO^ mice. A) Depletion of Gr1+ cells leads to mortality by 2 dpi. B) TLR4^KO^ animals have reduced Gr1+ cell infiltration at 4 hpi in the liver compared to control heterozygous littermates. C) TLR4^KO^ animals have reduced serum levels of Cxcl1, Cxcl10, Il-1β, and TNFIZl compared to control heterozygous littermates. D) TLR4^KO^ mice are resistant to abscess formation but have elevated CFUs relative to B6J controls that lack abscesses. E) BALB/cJ mice have reduced Gr1+ cell infiltration at 4 hpi compared to B6J and B6N. F) BALB/cJ mice have reduced serum levels of Cxcl1, Cxcl10, Il-1β, and TNFIZl compared to B6J and B6N mice, which have similarly reduced Gr1+ cell recruitment and serum cytokines. G) Similar *E. coli* hepatic CFU burden 4 hpi in BALB/cJ, B6J, and B6N mice.

Since BALB/cJ and TLR4^KO^ B6J mice were both resistant to abscess formation (Figure 3A, 7D), we assessed whether both strains share similarly reduced immune responses compared with wild-type B6J mice at 4 hpi. BALB/cJ mice had reduced Gr1^+^ immune cell infiltration, comparable to levels observed in TLR4^KO^ mice (Figure 7E). Furthermore, similar low serum levels of Cxcl1, Cxcl10, Il1β, and Tnfα were found in BALB/cJ and TLR4^KO^ B6J mice (Figure 7C and F). Importantly, at 4hpi, no difference in *E. coli* CFU burden was observed between BALB/cJ, B6J, and B6N mice, confirming that the differential immune response is induced prior to abscess-distinguishing replication (Figure 7G). Thus, at 4 hpi, two abscess-resistant strains of mice (BALB/cJ and TLR4^KO^) display similarly attenuated immune responses relative to susceptible mice (B6J and B6N). Together, these data suggest that following inoculation, *E. coli* in the liver signals via TLR4 to promote influx of innate immune cells, which in turn facilitates bacterial replication and development of liver abscesses. Although BALB/cJ mice possess functional TLR4, their early hepatic inflammatory response is likely blunted through other mechanisms.

### TLR4 governs a tradeoff between efficient clearance and abscess formation

B6J females lacking TLR4 had elevated hepatic bacterial burdens compared to WT B6J or TLR4^Het^ littermates that did not form abscesses (Fig 7D). The increase in CFU in TLR4^KO^ animals could be explained by a reduced capacity to clear the inoculum, to control bacterial replication, and/or to control pathogen dissemination between organs. To quantify the extent to which clearance, replication, or dissemination contribute to the increase in burden in TLR4^KO^ animals, we sequenced the barcode loci in *E. coli* and performed STAMPR analysis. This computational framework quantifies the number of cells from the inoculum that give rise to the population in an organ, known as the founding population (FP). Founders represent the organisms that survived infection bottlenecks, which consist of host factors that eliminate bacteria from the inoculum. A decrease in FP signifies a tightening of the infection bottleneck, and thus an increase in host clearance of the pathogen. The ratio of CFU to FP quantifies the net expansion of each clone; high CFU/FP ratios signify that each founding clone is represented multiple times, which in the absence of substantial dissemination, is due to bacterial replication. Finally, comparison of barcode frequencies between organs yields a genetic distance (GD) metric, where lower GD values indicate increased similarity between samples and therefore suggest increased dissemination.

TLR4^KO^ mice had higher founding populations (Figure 8B) compared to TLR4^Het^ littermates, indicating that TLR4 is required for efficient clearance of the inoculum. TLR4^Het^ animals that developed abscesses had substantially higher CFU/FP ratios than both TLR4^KO^ and TLR4^Het^ animals that did not develop abscesses (Figure 8C). However, when TLR4^Het^ animals that developed abscesses were excluded from the analysis, we found that the TLR4^KO^ mice possessed higher CFU/FP ratios; each clone was more abundant in TLR4^KO^ mice compared to heterozygote littermate (Figure 8C). The higher CFU/FP ratio in TLR4^KO^ mice is driven by increased bacterial replication, since neither TLR4^KO^ and TLR4^Het^ mice substantially shared bacteria between the liver and spleen (Figure 8D). These data together reveal that the increase in CFU in TLR4^KO^ animals is primarily due to a failure of the TLR4 deficient animals to clear the inoculum and their inability to control a subsequent ∼1-3 *E. coli* net cell divisions, but not due to an increase in dissemination. In contrast, TLR4^Het^ animals efficiently clear the inoculum, but surviving *E. coli* clones are more likely to undergo a net of ∼15-20 cell divisions within abscesses. Therefore, TLR4 signaling mediates a tradeoff between liver abscess development and efficient pathogen elimination.

**Figure 8.**
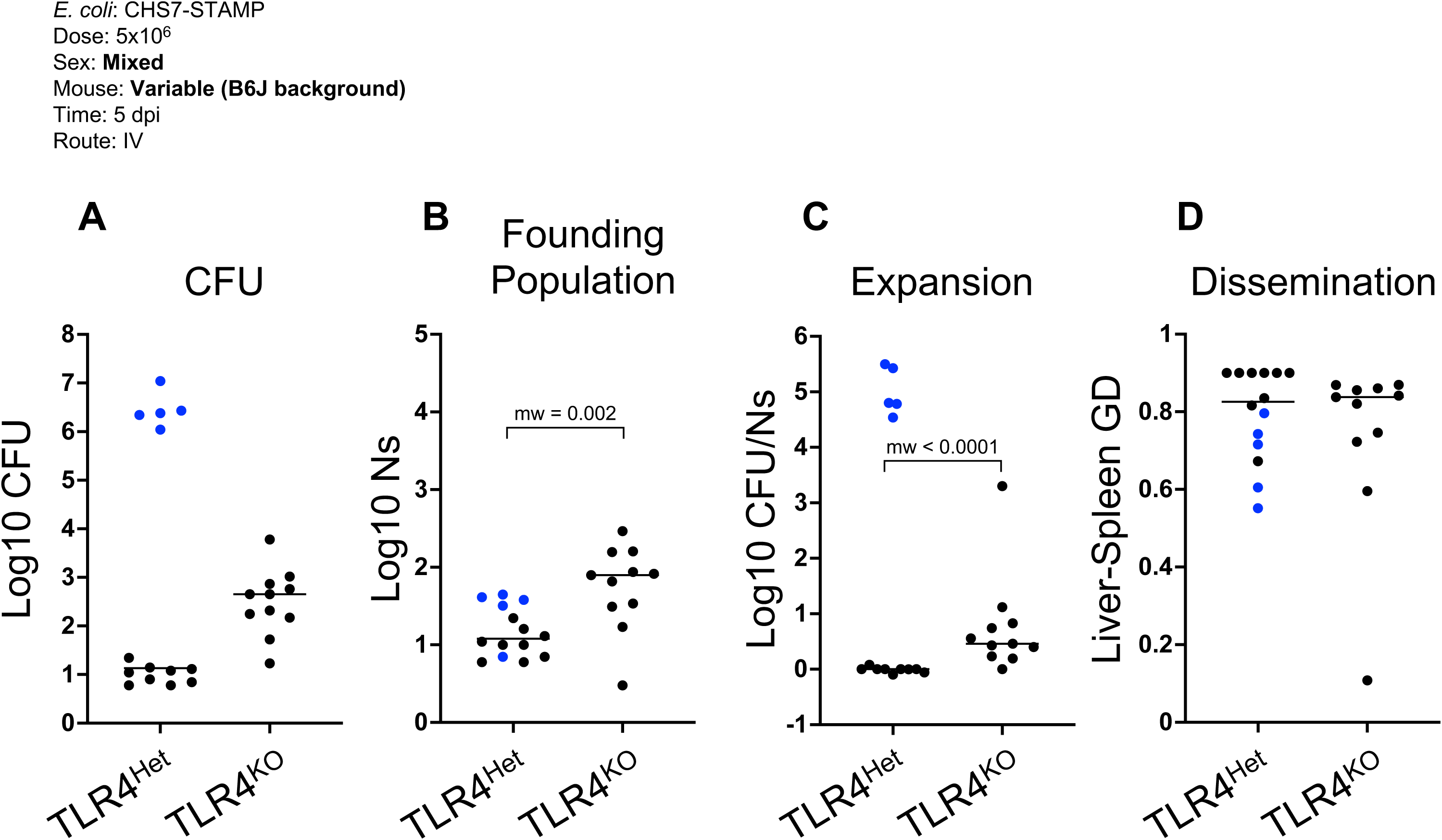
TLR4 controls infection bottlenecks and limits *E. coli* replication. A) E. coli hepatic CFU burden from TLR4^Het^ and TLR4^KO^ (as shown in Figure 7D but represented per ¼ liver for appropriate comparisons to founding population sizes). B) TLR4^KO^ mice have higher *E. coli* founding population sizes compared to TLR4^Het^. C) Measurement of net bacterial expansion (CFU per founder) indicates that abscesses (blue) contain a markedly expanded *E. coli* population. *E. coli* in TLR4^KO^ animals undergo more expansion compared to littermate controls that fail to form abscesses (black). D) Similar genetic distances between liver and spleen *E. coli* populations suggest that systemic dissemination is minimal and not influenced by TLR4.

## Discussion

Our study identifies molecular and genetic factors that govern tissue-specific liver abscess formation in a mouse model of *E. coli* systemic infection. In contrast to several other abscess and bacteremia models, mice with *E. coli* liver abscesses ultimately clear the infection, suggesting that this system will be valuable for understanding mechanisms that lead to abscess clearance as well as formation. Abscesses represent localized regions of hepatic necrosis and marked replication of relatively few *E. coli* clones (14). Since commensal *E. coli* can elicit abscess formation and TLR4^KO^ animals do not develop abscesses, we propose that abscesses result from exuberant TLR4-driven immune responses, rather that specific pathogen-derived virulence factors, which drive *Staphylococcus aureus*-induced renal abscesses (26). Further supporting the hypothesis that host factors primarily drive *E. coli* liver abscess formation is our observation that sex and mouse genotype govern abscess susceptibility. Relative to susceptible animals, mice that are resistant to abscess formation exhibit reduced Gr1+ immune cell recruitment and reduced proinflammatory cytokine production in the hours following infection, suggesting that early hepatic responses to *E. coli* may determine whether abscesses form. One to three days following inoculation, heterogeneous inflammatory immune cell clusters form in the liver, coinciding with the replication of a small number of *E. coli* clones. In the absence of Gr1^+^ cells, mice succumb to infection before abscesses can develop. However, mice that recruit fewer Gr1^+^ (BALB/cJ and TLR4^KO^) do not form abscesses and do not exhibit pathogen clonal expansion. Together our findings suggest that abscesses result from collateral damage caused by infiltrating innate immune cells and their products and that *E. coli* exploits areas of damaged tissue to replicate more substantially than in the absence of tissue damage (Figure S10). Control of the delicate balance between recruitment of sufficient inflammatory cells to abrogate *E. coli* replication and limiting damage to normal liver tissue by the inflammatory process appears to be defective in the livers of the C57BL/6 lineage. We speculate that defects in the mechanisms that govern this balance may underlie tissue-specific damage associated with a variety of infections.

Spatial transcriptomics is a powerful emerging approach for mapping the distribution and function of host cells in intact tissue, but to date has had limited application to infection contexts (27). At three and five dpi, we found that abscess cores were marked by a paucity of RNA signal and the absence of hepatic zonation markers, likely indicative of necrosis. Immune cell clusters adjacent to necrotic hepatocytes within abscess were highly heterogenous, consisting of dendritic cells, particularly *Cacnb3+* migratory dendritic cells, and other *Itgam*+ (CD11b) cells, innate lymphoid cells, T cells, neutrophils, monocytes, and macrophages (Figure 3, Figure S4). These immune clusters were enriched for markers of inflammatory responses such as lysozyme (*Lyz2*) that could contribute to tissue destruction. Unexpectedly, we found a rim of live hepatocytes expressing *Cxcl1* surrounding the abscesses by 5 dpi (Figure S4). Analyzing the specific functions of these cells, and how their roles in abscess formation and resolution is regulated by Cxcl1, should be experimentally approachable using CRISPR-based technology to genetically modify hepatocytes in vivo (28)

Single-cell RNA sequencing revealed a dramatic influx of macrophages and neutrophils into the liver four hpi. These two cell types, typically defined by expression of Ly6g (neutrophils) and Adgre1 (F4/80, macrophages), expressed a similar transcriptional program following infection, including *S100a8/9*, *Lcn2*, *Cxcl2*, and *Il1b* (Figure 6, Figure S8). Therefore, although infection induces infiltration of distinct cell lineages, they express similar genes and may play similar roles in pathogen clearance or abscess formation. We also found that changes in the abundance of immune cell populations and their respective transcriptional outputs was highly responsive to the *E. coli* dose. For example, a 5×10^4^ dose yielded a larger population of monocytes (cluster 5) relative to macrophages (cluster 1 and 3) (Figure 6), suggesting that higher doses, where a higher macrophage to monocyte ratio was observed, may lead to more efficient macrophage maturation and/or influx. The relative abundances of distinct cell types within the liver and their transcriptional states may be consequential for scaling the clearance capacity with the inoculum size. Distinct components of the innate immune response also scale with dose at different rates. For example, *Lcn2* and *Il1b* transcripts were induced within infiltrating cells at the lowest dose tested, but expression of *Cxcl1* and *Ifng* scaled more gradually and within distinct cell types (Figure S8). Together these observations uncover the key role of infectious dose in control of consequential immune responses, which likely modulate infection outcomes (22).

We found that abscess susceptibility depends on sex but is not directly linked to sex chromosomes. B6N females were more susceptible than B6N males, and F1 heterozygous females, from crosses between sensitive and resistant strains, are more susceptible than F1 heterozygous males. Given that *E. coli* abscess susceptibility is not directly due to alleles on the X, Y, or mitochondrial chromosome, we speculate that hormonal differences control expression of autosomal genes that confer abscess susceptibility. Sex bias has also been observed in *Entamoeba histolytica* liver abscesses in mice (29–32) and humans (33), where males are more susceptible to liver abscesses than females. In the mouse model, parasites are injected intrahepatically, and orchiectomy reduces abscess formation in males, suggesting a critical role for androgens in abscess susceptibility (30). Elucidating the mechanistic linkages between sex and abscess formation may have broader ramifications for deepening understanding of the well-documented sex differences in human immunity, such as the female bias for autoimmune disorders (34).

In our proposed model for abscess formation, following IV inoculation, *E. coli* that lodges in the liver stimulates recruitment of innate inflammatory cells that can provoke damage to the hepatic parenchyma, which in turn facilitates replication of *E. coli* clones (Figure S10). Bacterial replication then leads to recruitment of additional immune cells through positive feedback mechanisms, ultimately leading to abscess formation. Within the framework of our model, the apparent stochasticity in abscess formation in B6J animals can be explained by the observations that B6J mice appear to lie in the phenotypic space in-between resistant (BALB/cJ) and susceptible (B6N) animals; e.g., in CFU and number of replicating clones at 1 dpi, before abscesses have fully formed. Furthermore, we also observed that animals that are more likely to develop abscesses are also more likely to have higher CFU in livers containing abscesses, suggesting that the processes that control likelihood of abscess formation (frequency) may be intertwined with those that control their development (CFU) (Figure S9). Together, these observations suggest that the bimodality in B6J mice is driven by normally distributed immune responses that give rise to zero (resistant) or at least one (susceptible) replicating clones. Therefore, although mice that develop abscesses possess 100,000 times more CFU than mice that do not, the early immunologic events that appear to account for abscess formation may only differ subtly between susceptible and resistant mice, especially if they are the same genotype. We speculate that heightened abscess susceptibility in B6N mice is due to an increase in collateral damage caused by infiltrating cells, which facilitates the replication of a greater number of clones. Notably, the increase in collateral damage does not appear to cause sepsis or other negative clinical outcomes in B6N mice.

Our study adds *E. coli* to the few bacteria, including *Klebsiella pneumoniae* (35, 36) and *S. aureus* (37–39), that are known to give rise to large macroscopic liver abscesses in mice. However, the pathogenesis of the abscesses caused by these three pathogens appears to differ substantially. In marked contrast to the *E. coli* abscess model, mice that develop *K. pneumoniae* abscesses also succumb to infection and have high bacterial burden in other tissues (36, 40), suggesting that pathogen specific virulence factors, such as capsular polysaccharides, are sufficient to counter host defenses in a variety of tissues (41). *S. aureus* liver abscesses arise in humanized transgenic mice expressing HLA-DR4, owing to the direct stimulation of T cells by bacterial superantigens (37). Furthermore, *S. aureus* can form abscesses in the kidneys and skin even in wild-type strains of mice (42–44). In humans, *E. coli* is among the most common bacteria found within liver abscesses (15, 45–47), and understanding the molecular determinants of abscess formation and resolution in the murine model presented here may offer important insights for controlling human infections. Taken together, our study demonstrates that murine *E. coli* liver abscess provide a unique opportunity to decipher liver-specific innate immune mechanisms.

## Methods

### Ethics

All animal experiments were conducted in accordance with the recommendations in the Guide for the Care and Use of Laboratory Animals of the National Institutes of Health and the Animal Welfare Act of the United States Department of Agriculture using protocols reviewed and approved by Brigham and Women’s Hospital Committee on Animals (Institutional Animal Care and Use Committee (IACUC) protocol number 2016N000416 and Animal Welfare Assurance of Compliance number A4752-01).

### Animal experiments

8–12-week-old mice were used for all experiments. Both sexes are used in this study where indicated. Vendor-acquired mice were C57BL/6J (B6J, The Jackson Laboratory 000664), C57BL/6NJ (B6N, The Jackson Laboratory 005304), CBA/J (The Jackson Laboratory 000656), C3H/HeJ (The Jackson Laboratory 000659), BALB/cJ (The Jackson Laboratory 000651), CB6F1/J (The Jackson Laboratory 100007), and B6(Cg)-Tlr4tm1.2Karp/J (TLR4^KO^, The Jackson Laboratory 029015). Other F1 hybrids (BALB/cJ x B6J, BALB/cJ x B6N) and TLR4^KO^/TLR4^Het^ were bred at Brigham and Women’s Hospital. Animals were maintained at 68-75°C with 50% humidity in 12 hour day-night cycles.

For infections, defined volumes of frozen CHS7-STAMP library (14), Nissle, or MG1655 derived from overnight cultures were thawed, diluted in PBS, and immediately used to inoculate mice. For intravenous injections, animals were restrained using a Broome-style restrainer (Plas-Labs) and inoculated via the lateral tail vein with 100µl using a 27G needle. A heating pad was used to facilitate dilation of the tail vein. For intraperitoneal injections, animals were inoculated with 100µl into the abdominal cavity with a 27G needle. Gr-1 antibody (ThermoFisher 14-5931-85) was administered (100µg dose) by intraperitoneal injection 1 day prior to inoculation. At indicated times, animals were euthanized by isoflurane inhalation and cervical dislocation or cardiac bleed (when appropriate). To quantify CFU, organs were excised and homogenized with 2 x 2.3 mm stainless steel beads for 2 minutes with a bead beater (BioSpec). Organs were plated and diluted on LB + Kanamycin (for CHS7) or LB (for MG1655 and Nissle) plates.

### STAMPR analysis

Analysis of barcode frequency was performed as previously described (13, 14). Liver homogenates were plated as lawns and bacteria were scraped and diluted in PBS+25% glycerol and stored at -80°C. To amplify the barcode locus, samples were thawed and diluted in water and used as template for PCR (25 cycles). Amplicons were verified by agarose gel electrophoresis, pooled, column purified (GeneJet PCR Purification Kit), and sequenced on a MiSeq (Illumina) as 1×78 nt reads. Reads were trimmed and mapped to a defined list of barcodes in CLC Genomics Workbench (Qiagen). Read counts were exported and custom R scripts were used to visualize barcode frequencies and calculate founding population and genetic distance. In Figure 8, all FP and CFUs are reported for ¼ of the liver, which were homogenized in a total of 4 ml but only 1 ml was plated and scraped for STAMPR analysis.

### Backcross experiment and analysis

CB6F1 females were crossed with C57Bl/6J males. On weaning, male offspring were genotyped using the Transnetyx Genetic Monitoring service. Male N1s were infected at 9-12 weeks of age and CFU and abscess formation was assessed in the liver at 5 days post infection. Of the ∼10,000 SNPs that are genotyped, ∼3,000 distinguish BALB/cJ and B6J. Genotyping data was first converted to binary heterozygous (0) or homozygous (1) calls. Since there is only one copy of the X chromosome in male N1s, the BALB/cJ allele was treated as heterozygous (0), and the B6J allele was treated as homozygous (1). For every SNP, mice were separated into homozygous or heterozygous bins, and the abscess frequency within each bin was calculated. The abscess frequency in the homozygous bin relative to the abscess frequency in the heterozygous bin is the Y-axis in Figure S6.

To validate the binning approach described above, animals were also monitored for coat color, which is governed by the agouti locus in Chromosome 2; in N1 males, heterozygous mice have brown coats, and homozygous mice have black coats. We separated mice at every SNP into bins as described above and calculated “black coat frequency” in each group. As expected, when binning near the agouti locus in Chromosome 2, 100% of mice in the homozygous bin have black coats, and 0% of mice in the heterozygous bin have black coats, confirming the validity of our approach at detecting monogenic traits. To avoid dividing by 0, we assume that 0.5 mice in the heterozygous bin had a black coat, yielding a log2 fold change of ∼8. Importantly, a 50% penetrant monogenic trait would be expected to have a log2 fold change of ∼7 with our sample size (153 mice). Since abscesses are ∼70% penetrant in inbred B6J males, a peak would have been evident if the trait was monogenic.

### Flow cytometry

To obtain liver cell suspensions, livers were excised and minced with scissors in HBSS + 10 mM EDTA in a 50 ml conical tube. Tissue was then washed 3x with 50 ml PBS to remove EDTA. After the tissue settled to the bottom of the tube, PBS was carefully removed and replaced with 10ml DMEM containing 0.2 mg/ml DNase (Roche 10104159001) and 1 mg/ml Collagenase (Sigma-Aldrich C5138). Tissue was then incubated for 30 minutes at 37°C and passed through a 70µm filter. Additional DMEM washes and mechanical force with a syringe plunger were used to propel cells stuck on the filter through. Cells were centrifuged at 50xg for 5 minutes to spin down hepatocytes, and supernatants, enriched for nonparenchymal cells, were placed in a new 50 ml conical tube. These cells were centrifuged at 500xg for 5 minutes, washed with 10ml of PBS, transferred to a 15 ml conical tube, and centrifuged at 500xg for 5 minutes. The supernatant was removed and 1 ml of red blood cell lysis buffer (Roche 11814389001) was added and cells were incubated for 1 minute, after which 10 ml of PBS was added. Cells were centrifuged at 500xg for 5 minutes and resuspended in 2ml of PBS. To prepare a Percoll gradient, a long Pasteur pipette was used to introduce Percoll to the bottom of the cell suspension. 2ml of 40% Percoll (prepared in HBSS and diluted in DMEM) was added, followed by 2 ml of 80% Percoll. The gradient was centrifuged for 1300xg for 20 minutes. Cells between the 80% and 40% layers were carefully removed and washed in 10 ml of PBS. The cells were then resuspended in 1ml of PBS with 2mM EDTA and 2% FBS. Antibodies (anti-CD45 [Biolegend 103129] and anti-Gr1 [Invitrogen 53-5931-82]) were added to cell suspensions at 1:200 dilutions and incubated at 4°C for 30 minutes. Cells were centrifuged and resuspended in 200µl of PBS with 2mM EDTA and 2% FBS. Flow cytometry was performed with an SH100 Cell Sorter (Sony Biotech) and analyzed with FlowJo.

### Cytokine and ALT measurements

Blood was collected via cardiac bleed and left to coagulate in 1.5 ml tubes at room temperature. Serum was retrieved following centrifugation at 2,000xg for 10 minutes at 4°C. ALT was measured with the Alanine Transaminase Colorimetric Activity Assay Kit (Cayman Chemical 700260) according to the manufacturer’s instructions. Cytokines were measured by multiplexed bead-based protein capture (EveTechnologies)

### Histology

Livers were embedded in a 30% sucrose:OCT (1:2.5) solution, frozen immediately, and stored at -80°C. Hematoxylin and eosin staining was performed at the Harvard Rodent Histopathology Core facility. Slides were imaged with an Eclipse Ti microscope.

### Single-cell RNA sequencing

Mice were infected as described above and euthanized 4 hours post inoculation. Livers were processed as above for flow cytometry, but without DNAse and Percoll to minimize preparation time. Cells were sorted by CD45 expression into PBS + 2% FBS. Cells were processed using the Chromium Next GEM Single Cell 3’ Reagent Kits (10x genomics) and sequenced on a NovaSeq 6000 (Illumina) at the Harvard Medical School Biopolymers Core Facility as 28 (read 1) and 90 (read 2) nt reads.

Reads were processed with 10X Genomics Cloud Analysis to generate hdf5 files and further analysis was performed with Seurat v4.3 (48). Data were filtered by nFeature_RNA > 200, nCount_RNA > 1000, and percent.mt < 80 and normalized with SCTransform. RunPCA and RunUMAP were used prior to doublet removal with DoubletFinder (pN = 0.25, pK = 0.09). Data were the integrated with FindIntegrationAnchors and IntegrateData, after which PCA (RunPCA), cluster identification (FindNeighbors, dims = 1:15, and FindClusters), and UMAP (RunUMAP, reduction “pca”, n.neihbors = 20, min.dist = 0.3, spread = 1, metric = “Euclidean”) was performed. Data displayed in Figure 6 and Figure S8 are SCT transformed.

### Spatially resolved transcriptomics

Spatial transcriptomics were performed using MERSCOPE (Vizgen). Livers were embedded in a 30% sucrose:OCT (1:2.5) solution, frozen immediately, and stored at -80°C. Blocks were cut to 10µm sections with a CM1860 UV cryostat (Leica) on to MERSCOPE slides, which contain fluorescent beads for autofocusing on the MERSCOPE instrument. The slides were fixed in 4% paraformaldehyde in PBS (Fixation Buffer) for 15 minutes and washed 3x in PBS and then incubated with 70% ethanol at 4°C overnight.

Permeabilized sections were stained for cell boundaries with the Cell Boundary Staining Kit (Vizgen 10400009). Briefly, slides were washed once in PBS and incubated for 1 hour with Blocking Solution at room temperature. Slides were then incubated with Primary Staining Solution for 1 hour at room temperature. After 3 washes with PBS, slides were incubated with Secondary Staining Solution for 1 hour at room temperature. Slides were then washed 3x in PBS, incubated with Fixation Buffer for 15 minutes, and washed 2x with PBS. To hybridize probes, slides were first washed with Sample Prep Wash Buffer and incubated in Formamide Wash Buffer for 30 minutes at 37°C. 50µl of the MERSCOPE gene panel, a pre-defined panel that targets 140 genes (Table S1), was added to the tissue section, and incubated for two days at 37°C in a humidified chamber. Slides were then washed 2x for 30 minutes each in Formamide Wash Buffer at 47°C, and 1x in Sample Prep Wash Buffer.

Sections were then embedded in a thin gel consisting of a Gel Embedding Premix, 0.05% ammonium persulfate, and 0.005% N,N tetramethylethylenediamine. Gel-embedded slides were cleared by incubating in a Clearing Solution containing 1% Proteinase K at 37°C for three days at 37°C. Cleared sections were then imaged with the MERSCOPE instrument. Data was visualized and analyzed with the MERSCOPE Visualizer.

## Data Availability

Single Cell RNA Sequencing (PRJNA945406) reads have been deposited in the Sequencing Read Archive (SRA).

## Acknowledgements

This work is supported by NIH F31 AI156949 (K.H.), NIH R01 AI042347 (M.K.W.), and the Howard Hughes Medical Institute (M.K.W.). We are grateful to Caitlyn Holmes, Aric Brown, and members of the Waldor Lab for feedback on this manuscript, and to Jonathan Kagan for valuable discussion and advice. The Harvard Rodent Histopathology and Harvard Biopolymers core facilities supported this work.

## Figure Legends

**Figure S1.**
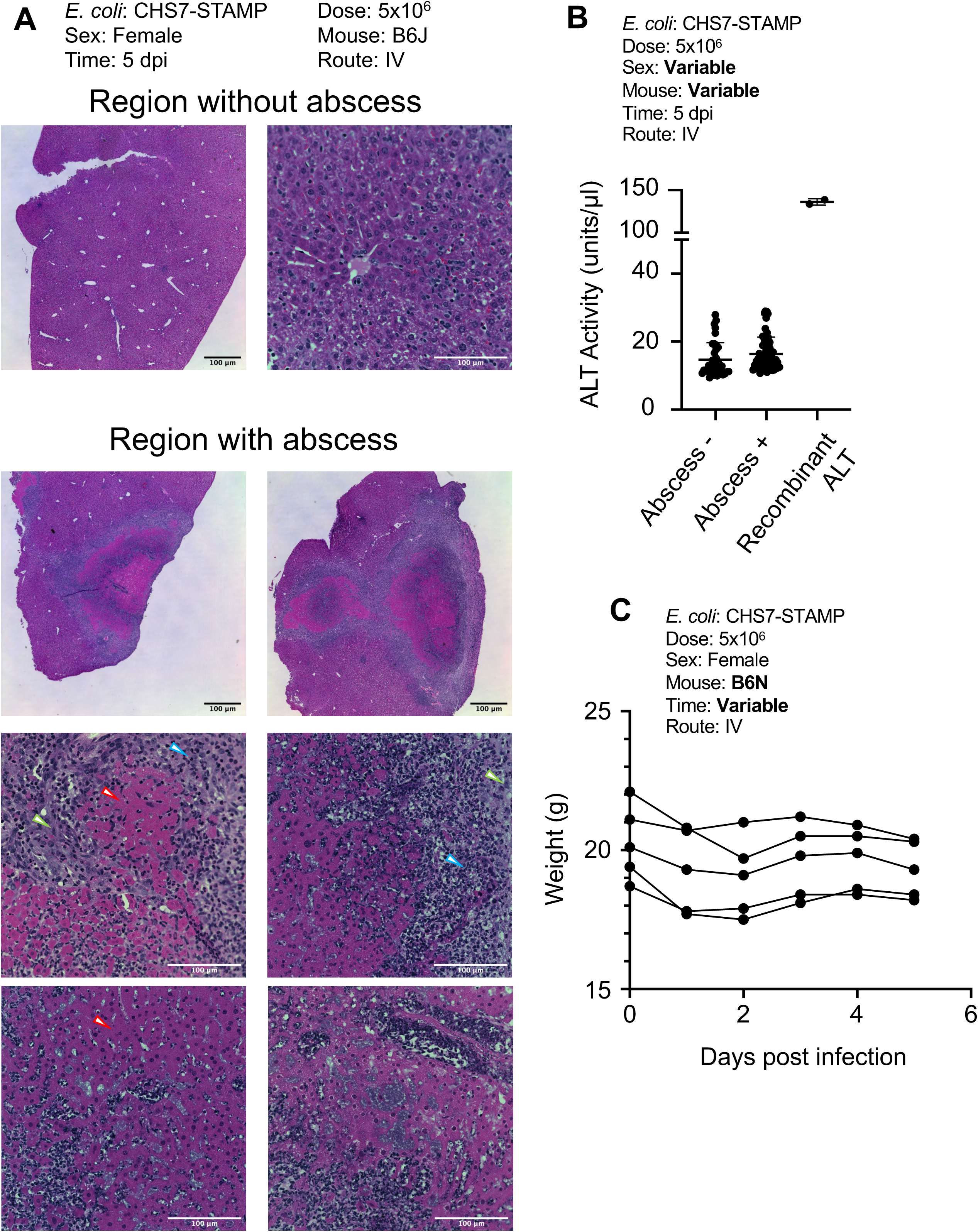
Histology and health outcomes of *E. coli*-induced liver abscesses. A) H&E staining of abscesses in B6J females. Necrotic hepatocytes (deep pink stain, red arrowheads) are surrounded by cells that resemble macrophages (green arrowhead) and neutrophils (blue arrowhead). B) Serum ALT levels are similar in mice that contain or lack abscesses. Serum was collected from mice used in Figure 2 and Figure 4. C) B6N female mice exhibit early weight loss up to 2 dpi and then stabilize in weight.

**Figure S2.**
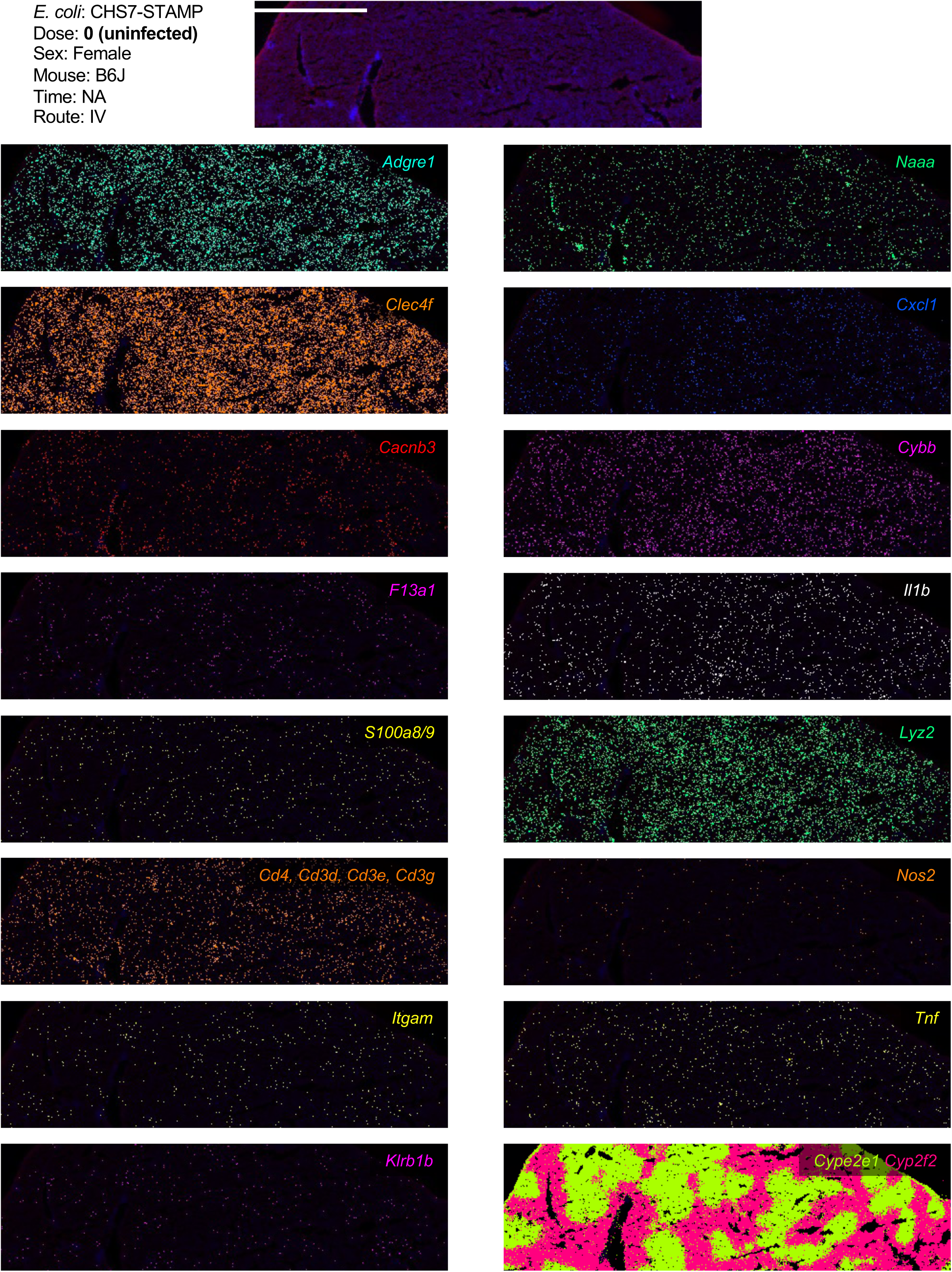
Spatial transcriptomic profile of uninfected liver tissue. MERSCOPE images of liver from uninfected samples from Figure 3 but individual genes are shown here separately. Cell types and corresponding transcripts are macrophages (*Adgre1),* migratory dendritic cells (*Cacnb3*), Kupffer cells (*Clec4f*), monocytes (*F13a1*), neutrophils (*S100a8/9*), T cells (*Cd4, Cd3g, Cd3e,* and *Cd3b*), various leukocytes (*Itgam*, also known as CD11b), NK/ILC1s (*Klrb1b*), and cDC1s (*Naaa*). *Cxcl1, Cybb, Il1b, Lyz2, Nos2,* and *Tnf* correspond to known markers of inflammatory responses. *Cyp2e1* and *Cyp2f2* correspond to markers of hepatocyte zonation.

**Figure S3.**
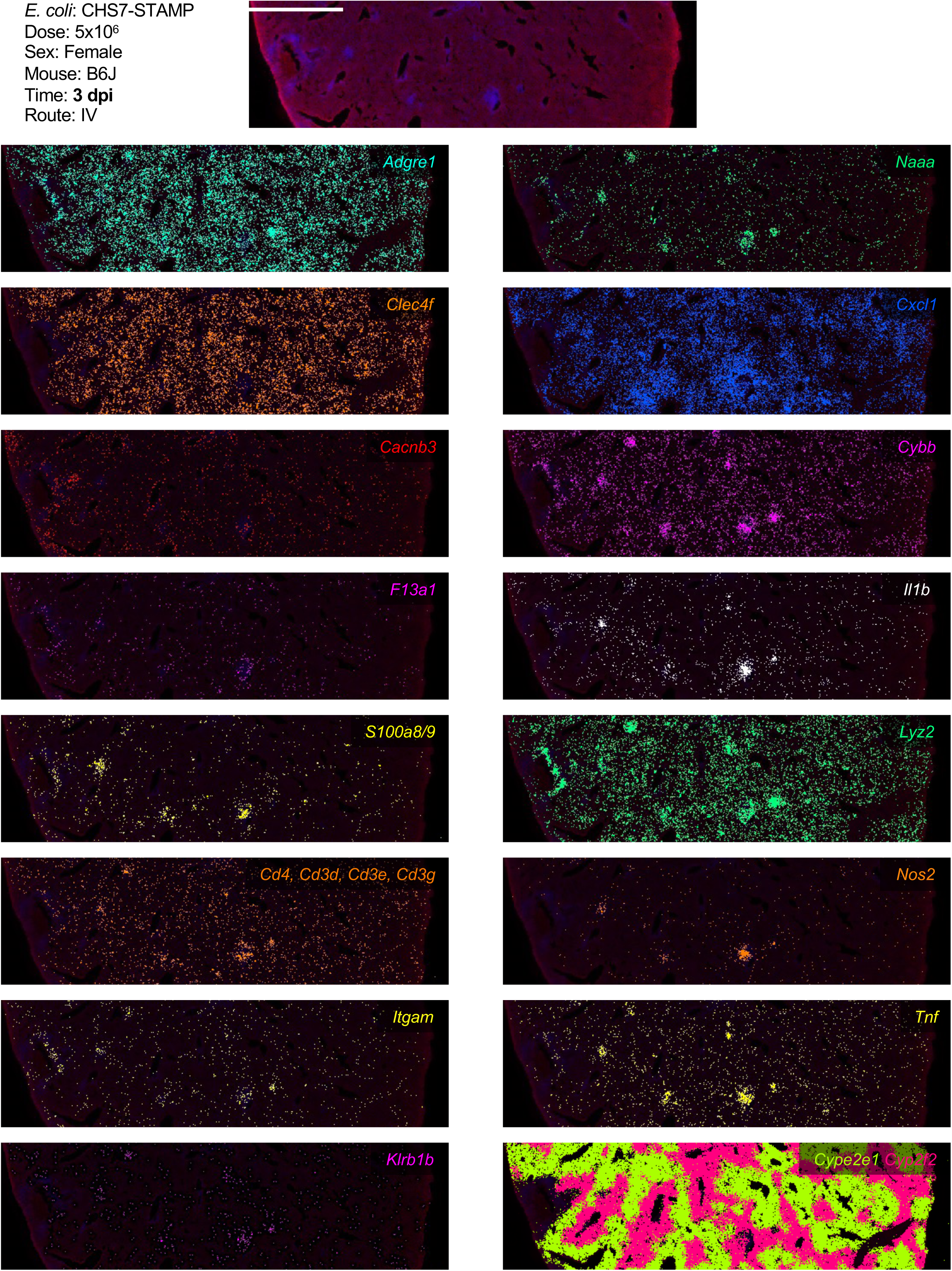
Spatial transcriptomic profile of 3 dpi abscess. MERSCOPE images of liver from 3 dpi samples from Figure 3 but individual genes are shown here separately. Cell types and corresponding transcripts are macrophages (*Adgre1),* migratory dendritic cells (*Cacnb3*), Kupffer cells (*Clec4f*), monocytes (*F13a1*), neutrophils (*S100a8/9*), T cells (*Cd4, Cd3g, Cd3e,* and *Cd3b*), various leukocytes (*Itgam*, also known as CD11b), NK/ILC1s (*Klrb1b*), and cDC1s (*Naaa*). *Cxcl1, Cybb, Il1b, Lyz2, Nos2,* and *Tnf* correspond to known markers of inflammatory responses. *Cyp2e1* and *Cyp2f2* correspond to markers of hepatocyte zonation.

**Figure S4.**
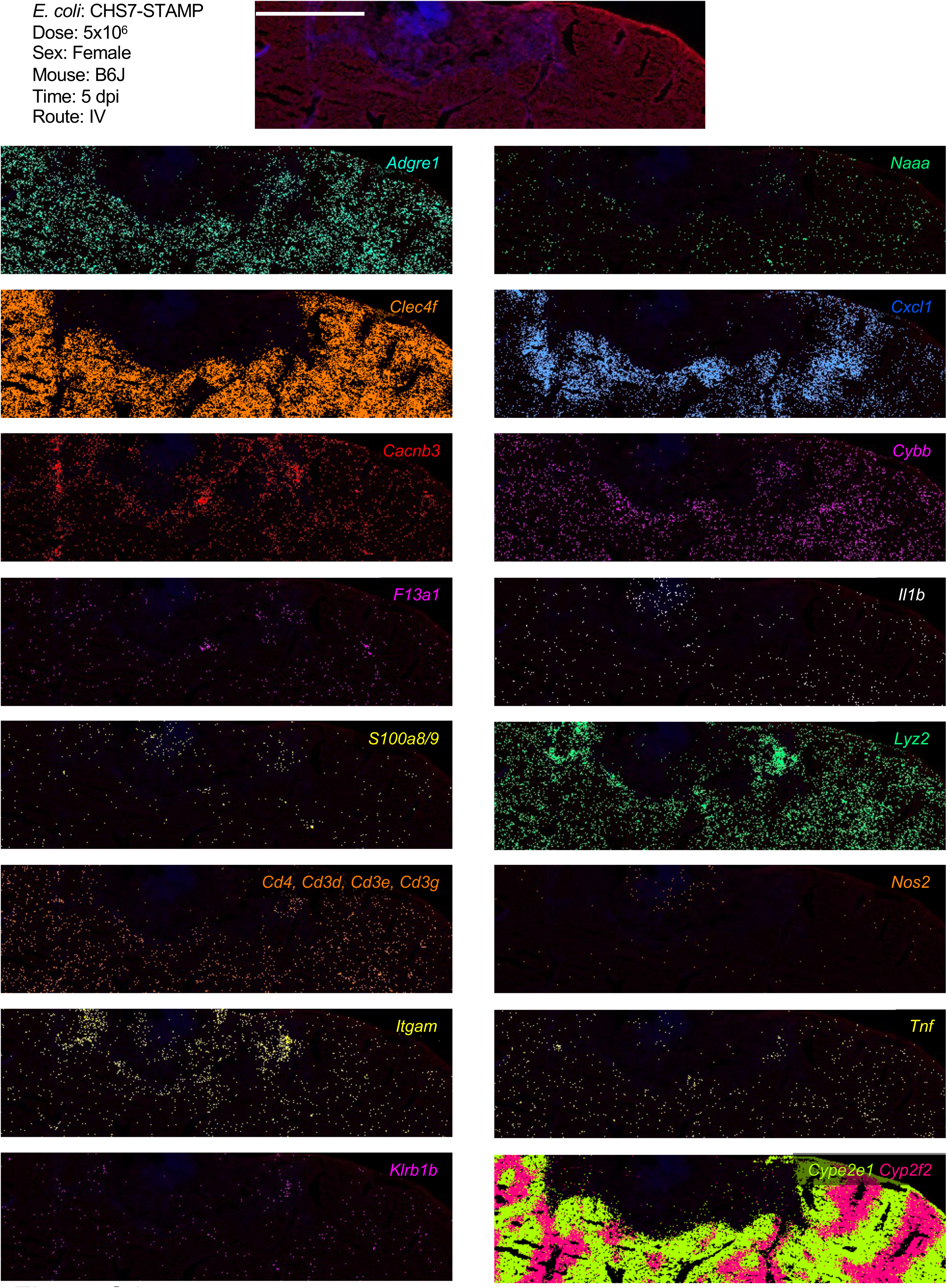
Spatial transcriptomic profile of 5 dpi abscess. MERSCOPE images of liver from 5 dpi samples from Figure 3 but individual genes are shown here separately. Cell types and corresponding transcripts are macrophages (*Adgre1),* migratory dendritic cells (*Cacnb3*), Kupffer cells (*Clec4f*), monocytes (*F13a1*), neutrophils (*S100a8/9*), T cells (*Cd4, Cd3g, Cd3e,* and *Cd3b*), various leukocytes (*Itgam*, also known as CD11b), NK/ILC1s (*Klrb1b*), and cDC1s (*Naaa*). *Cxcl1, Cybb, Il1b, Lyz2, Nos2,* and *Tnf* correspond to known markers of inflammatory responses. *Cyp2e1* and *Cyp2f2* correspond to markers of hepatocyte zonation.

**Figure S5.**
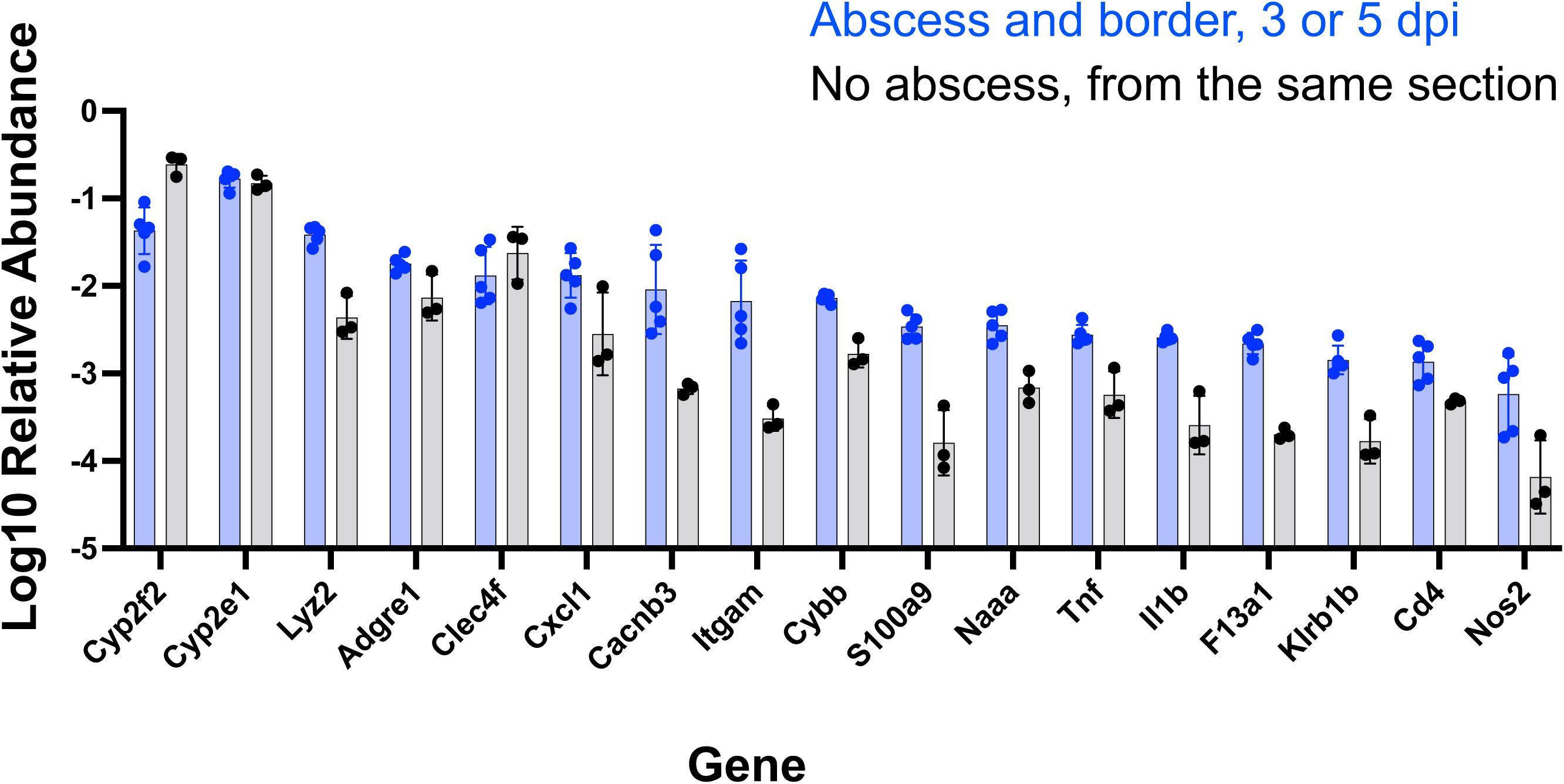
Quantification of MERSCOPE data. Quantification of relative transcript abundances within abscesses. Regions of interest were drawn around an abscess and bordering regions (blue) or control regions from the same section that lacked immune cell clusters (black). Transcripts were quantified relative to the total number of transcripts in the region. Data are derived from 3 and 5 dpi samples from 3 sections across two animals.

**Figure S6.**
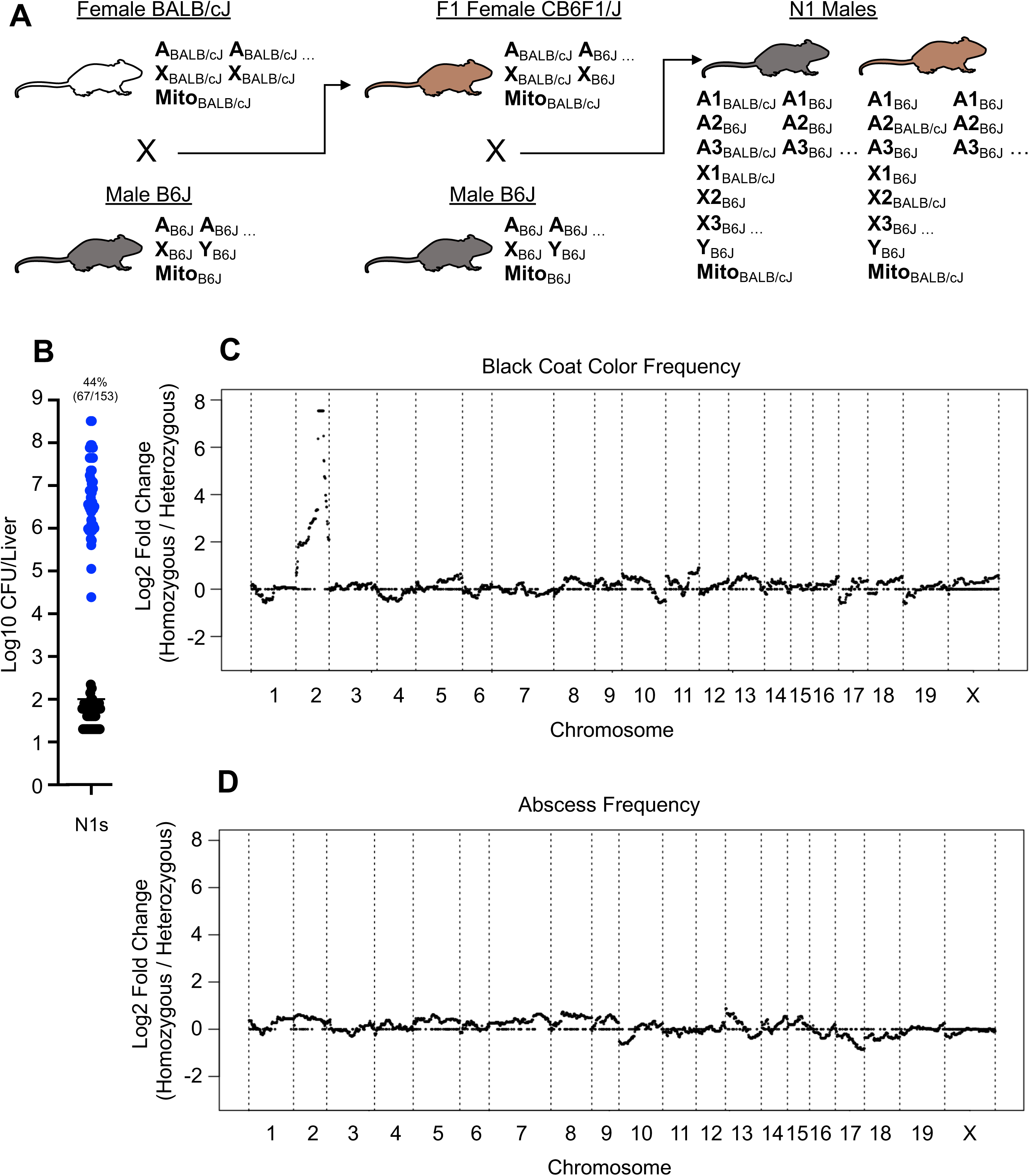
Backcross analysis of inheritance of abscess susceptibility. A) CB6F1/J heterozygotes were bred to male B6J mice to generate N1 backcross mice. The abscess-resistant phenotype of male heterozygous mice was expected to revert to susceptible after backcrossing with susceptible B6J mice. Therefore, only males that are homozygous B6J for the causal allele should develop abscesses. B) 44% of male N1 mice developed abscesses (blue). C) The agouti locus is identified when calculating the frequency of mice with black coat colors in homozygotes, relative to the frequency of mice with black coat color in heterozygotes. D) Same as C) but for abscess frequency instead of coat color. No association was observed between abscess susceptibility and B6J homozygosity.

**Figure S7.**
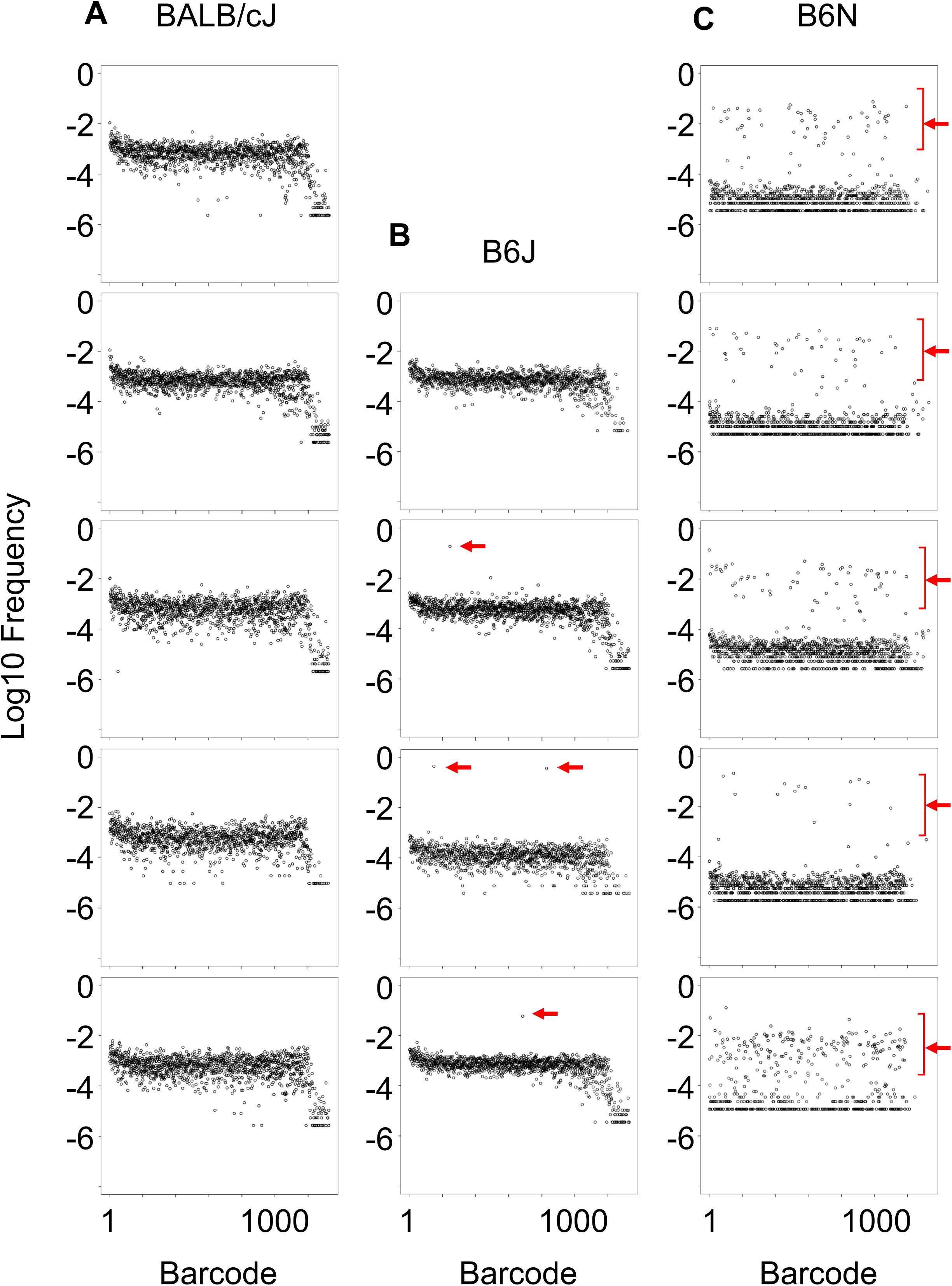
Clonal replication at 1 dpi correlates with abscess frequency. Data are replicate animals from Figure 5C. The X-axis is an arbitrary designation for barcode identity, and the Y-axis represents the relative frequency of each barcode. Red arrows denote replicated clones. A, B, and C correspond to BALB/cJ, B6J and B6N mice, respectively.

**Figure S8.**
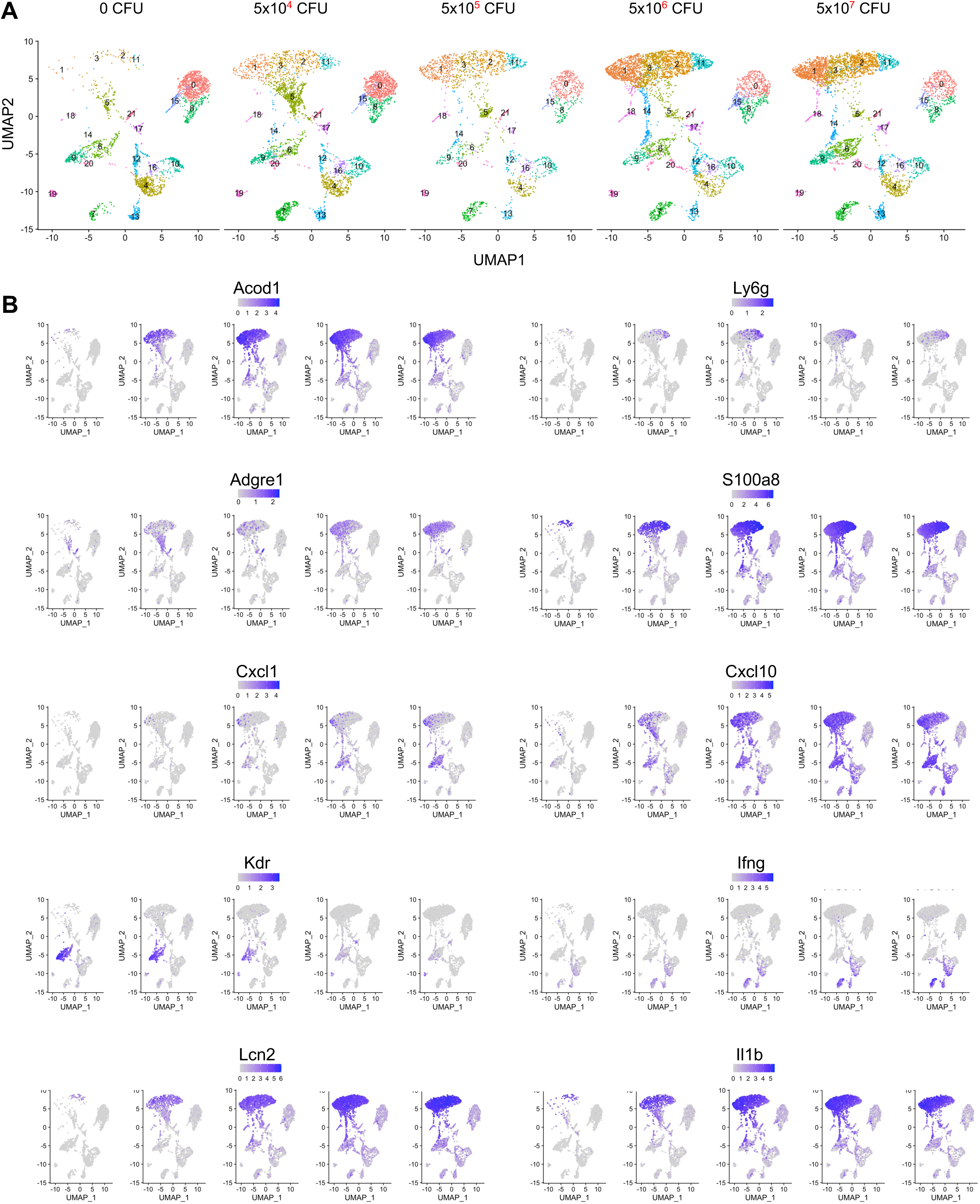
Additional genes from single cell RNA-sequencing of liver immune cells at 4hpi. A) UMAP plots are shown from Figure 6 for reference. B) Individual genes are shown as indicated from normalized expression data (sctransform).

**Figure S9.**
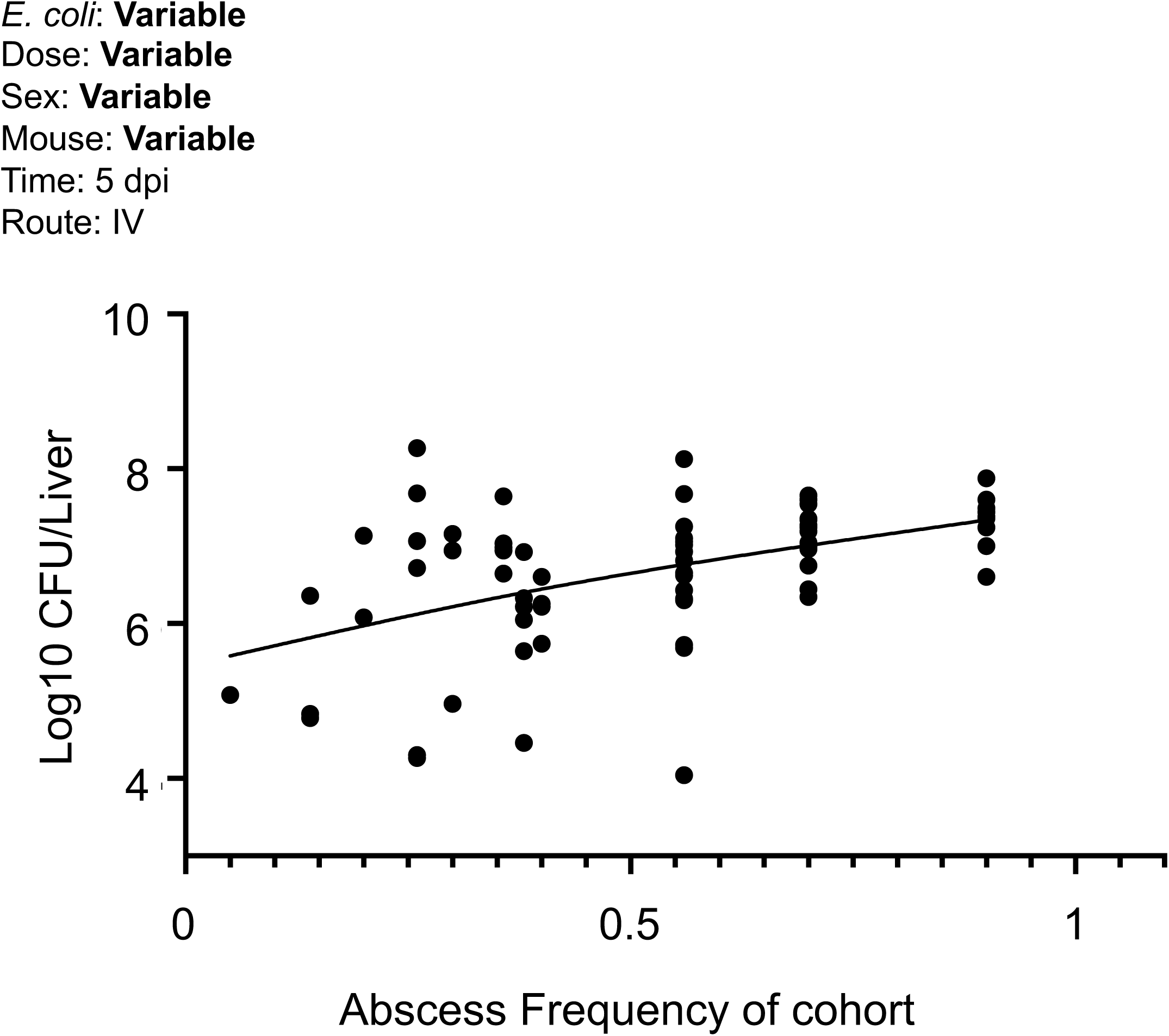
Abscess frequency is correlated with CFU. From every experiment in this study that assessed abscess frequency at 5 dpi, we plotted the CFU of animals that developed abscesses as a function of the frequency of the abscess within the experimental cohort. Both variables are positively correlated (Spearman r < 0.0001)

**Figure S10.**
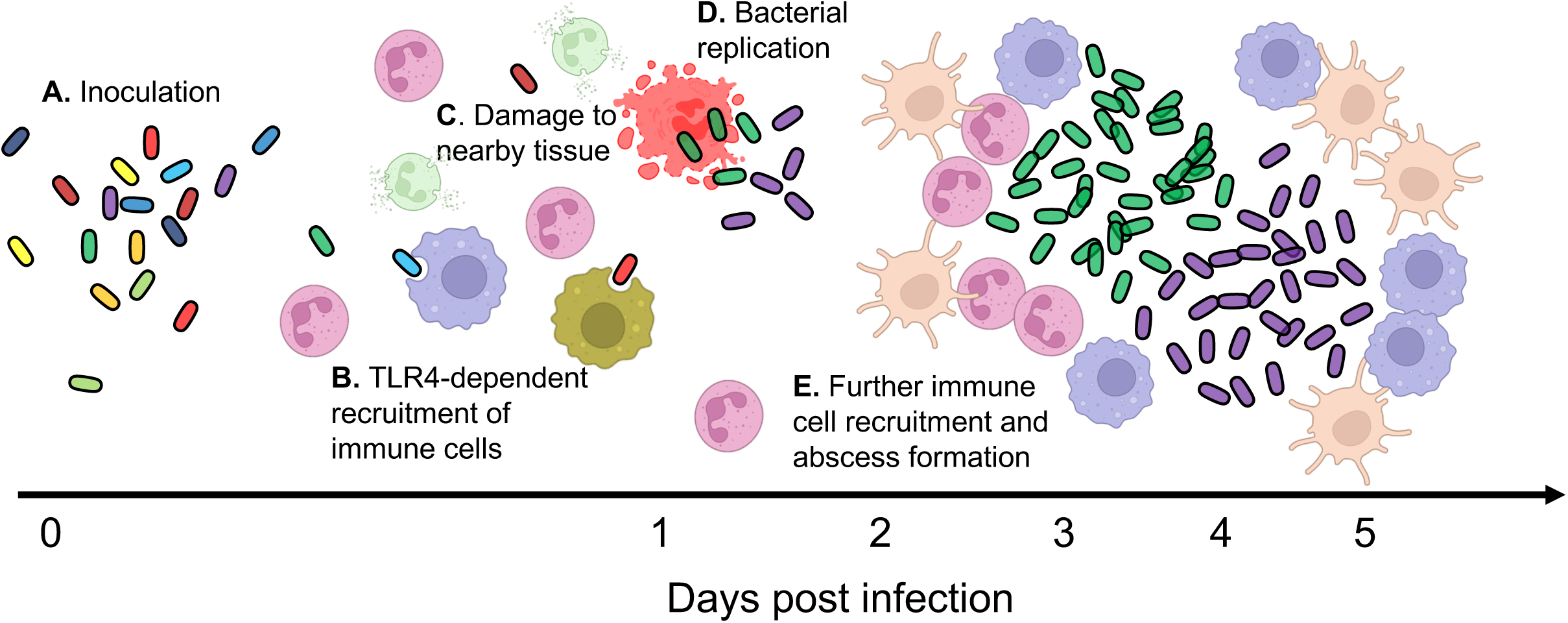
Proposed model for *E. coli* liver abscess formation. A) Inoculation of bacteria leads to the rapid recruitment of immune cells to the liver in a TLR4-dependent manner (B). Recruited inflammatory cells cause damage to neighboring tissue (C). *E. coli* exploits the newly necrotic niche to replicate by one day post inoculation (D), which leads to further recruitment of inflammatory cells and pathogen replication, until the abscess is fully formed (E).

